# Rapamycin Mitigates Branched-Chain Amino Acid Accumulation in Maple Syrup Urine Disease Mice

**DOI:** 10.64898/2026.11.10.765222

**Authors:** Swankita Godara, Morgan E. Schroeder, Isabelle A. Andsager, Ava P. Hickey, Raghbendra K. Dutta, Katie Cheng, Katherine E. Vest, Anthony S. Grillo

## Abstract

Maple syrup urine disease (MSUD) is an inborn error of metabolism caused by mutations in the mitochondrial branched-chain α-keto acid dehydrogenase (BCKDH) complex, resulting in toxic accumulation of branched-chain amino acids (BCAAs) and their corresponding branched-chain keto acids (BCKAs). This leads to poorly understood clinical features in both brain and muscle. Due to a limited understanding of MSUD pathophysiology, there is a marked paucity of effective pharmacological interventions. Here, we discovered the mTOR inhibitor, rapamycin, improves disease and extends survival in MSUD mouse models. Rapamycin treatment attenuates neuroinflammation and reduces muscle pathology. To investigate the mechanism behind these effects, we studied the accumulation of BCAAs and BCKAs. Rapamycin decreased BCAA accumulation. Overall, our work identifies rapamycin as a potential therapy for MSUD. More broadly, it suggests dysregulated mTOR signaling may drive MSUD pathobiology and advances our understanding of the mechanisms influencing leucine toxicity.

## INTRODUCTION

Inborn errors of metabolism (IEMs) represent diverse inherited diseases that can often lead to the accumulation of toxic metabolites, impair vital biochemical pathways, and manifest in a variety of symptoms affecting multiple organs such as brain and muscle^1–3^. Effective therapeutic options remain limited for >80% of these diseases affecting 1:2,500 births^3^. Notably, deficiencies in the proper metabolism of amino acids represent up to 50% of all IEM diagnoses from newborn screening cohorts^4,5^. Their multivariate and complex role in metabolism requires a deeper understanding of the molecular, biochemical, physiologic, and cellular processes that drive disease to develop rational targeted therapies^3^. Amino acid disorders are especially prevalent in IEMs^1–3^, and amino acid imbalance is also implicated in diabetes, cancer, liver disease, and normative aging^6–8^.

Maple Syrup Urine Disease (MSUD) is an IEM caused by the improper breakdown of the branched-chain amino acids (BCAAs) leucine, isoleucine, and valine, and their corresponding ketoacids (BCKAs)^9^. Deficiencies in the enzyme complex branched-chain α-keto acid dehydrogenase (BCKDH) preclude the irreversible oxidative decarboxylation of BCKAs to their CoA derivatives for energy production. MSUD is most often diagnosed as classic (cMSUD, ∼70% of cases) or as intermediate (iMSUD, ∼20% of cases) subtypes^10^ and is associated with ∼200 known BCKDH variants that lead to no or partial BCKDH activity, respectively^11^. MSUD patients exhibit symptoms including sweet-smelling urine, reduced growth, feeding difficulties, lethargy, developmental delays, respiratory distress, encephalopathy, neurologic complications (e.g. seizures), and skeletal muscle abnormalities including hypotonia, myofiber necrosis, and myofibrillar damage^11–16^.

MSUD pathophysiology remains poorly defined, but proposed mechanisms all largely converge on leucine cytotoxicity^9^. Elevated BCAAs may competitively inhibit uptake of neurotransmitter precursors through the large neutral amino acid transporter LAT1 (SLC7A5)^17^, reducing serotonin, dopamine, glutamate, or GABA production.^9^ This can impair neurologic signaling and muscle function, while reduced brain glutamate also disrupts neurotransmission and mitochondrial bioenergetics^18^. BCKAs, particularly α-ketoisocaproate (KIC), may further induce redox imbalance and neuroinflammation and impair mitochondrial oxygen consumption, membrane potential, and ATP/ADP balance^19^. Some patients exhibit leucine-sensitive hypoglycemia and muscle insulin resistance^12^ while >10-fold BCAA accumulation may disrupt proteostasis or promote unintended BCAA incorporation into proteins through mischarged tRNAs. Despite extensive study, there is no consensus on the mechanisms driving MSUD progression.

The absence of a clear mechanism has limited the development of effective pharmacological interventions. Early diagnosis and severe dietary leucine restriction using specialty formulas and medical foods improve disease prognosis^11,12,20^, but limit normal food intake and quality of life. This begins immediately after diagnosis and requires continued nutritional supplementation with frequent biochemical outpatient monitoring of plasma BCAA status. Suboptimal dietary nonadherence remains a major clinical problem^21^. Even patients who strictly adhere to dietary management remain vulnerable to metabolic crisis caused by infection, stress, or excessive leucine intake^21–23^. Liver transplantation greatly improves metabolic stability but is invasive, expensive, limited by organ availability, and potentially associated with long-term complications^20,24,25^. Cognitive dysfunction, impaired growth, ADHD, depression, and anxiety persist in MSUD patients^11,18^, and neither leucine restriction nor transplantation reliably restores cognitive function^18,26^. Thus, new approaches are needed.

Leucine is both a toxic metabolite in MSUD and a potent activator of mechanistic target of rapamycin (mTOR). mTOR plays a key role in the regulation of protein synthesis, autophagy, metabolism, growth, survival, and other major cellular processes with nutritional adaptation or in response to other environmental stimuli. The recent discoveries of leucine sensors, including Sestrin2^27^, SAR1B^28^, and LARS1^29^, have established direct molecular links between intracellular leucine availability and mTOR Complex 1 activation. Thus, leucine accumulation in MSUD may not only exert direct metabolic and neurotoxic effects, but also chronically perturb nutrient-sensing pathways. Consistent with this possibility, metformin reduces KIC accumulation in patient-derived MSUD fibroblasts^14^, and subsequent studies in MSUD models have implicated altered mTOR signaling and BCKA-dependent regulation of the Sestrin2–GATOR2 complex^30^. Previous work further demonstrated that the mTOR inhibitor rapamycin attenuates pathology and improves survival in severe mitochondrial disease^31,32^, and BCAA restriction is beneficial in normative aging by reducing mTOR activity^33,34^. These findings provided a rationale for testing whether mTOR inhibition could similarly modify MSUD progression.

Here, we show that rapamycin improves disease outcomes and extends survival in mouse models of MSUD, establishing this inborn error of metabolism as responsive to mTOR inhibition. Rapamycin attenuates neuroinflammation, neuropathological changes, and skeletal muscle degeneration. It further promotes beneficial therapeutic effects across the major tissues affected by MSUD. These improvements are accompanied by reduced accumulation of BCAAs and BCKAs in serum and disease-relevant tissues, associating physiological rescue to correction of the underlying metabolic disturbance. Together, our findings suggest maladaptation in mTOR signaling is a contributor to MSUD pathobiology and establishes rapamycin as a potential pharmacological intervention for this currently untreatable disease.

## RESULTS

### Rapamycin extends survival and suppresses disease progression in MSUD mouse models

Noting how leucine is a critical pathogenic BCAA in MSUD and activates mTOR, we hypothesized that mTOR activity drives disease progression in MSUD mouse models. We tested whether the potent mTOR inhibitor rapamycin alleviates MSUD symptoms in leading mouse models. Global knockout of the BCKDH E2 subunit (Dbt) produces cMSUD mice that model the null BCKDH mutations found in patients. These neonates exhibit >10-fold increases in serum, muscle, and brain BCAAs, excess BCKA accumulation, severe motor-cortex astrogliosis by GFAP staining, and encephalopathy causing death within 1-2 days after birth^10^. Whereas control cMSUD neonates died within two days, rapamycin beginning at PND0 (4 mg/kg/day, i.p.) increased median and maximal lifespan by >1,000% (**Fig. 1a**), comparable with genotype-agnostic benefits recently reported using mRNA therapy^35^. Consistent with the importance of mTOR in neonatal development, neonates treated with rapamycin were smaller than WT controls. However, these mice gained weight steadily (**Fig. 1b**) and survivors could be weaned around 30 days of age. Because the short lifespan of untreated cMSUD neonates precludes longitudinal physiological and behavioral studies, we next tested whether rapamycin similarly improves survival in other MSUD mouse models.

**Figure 1.**
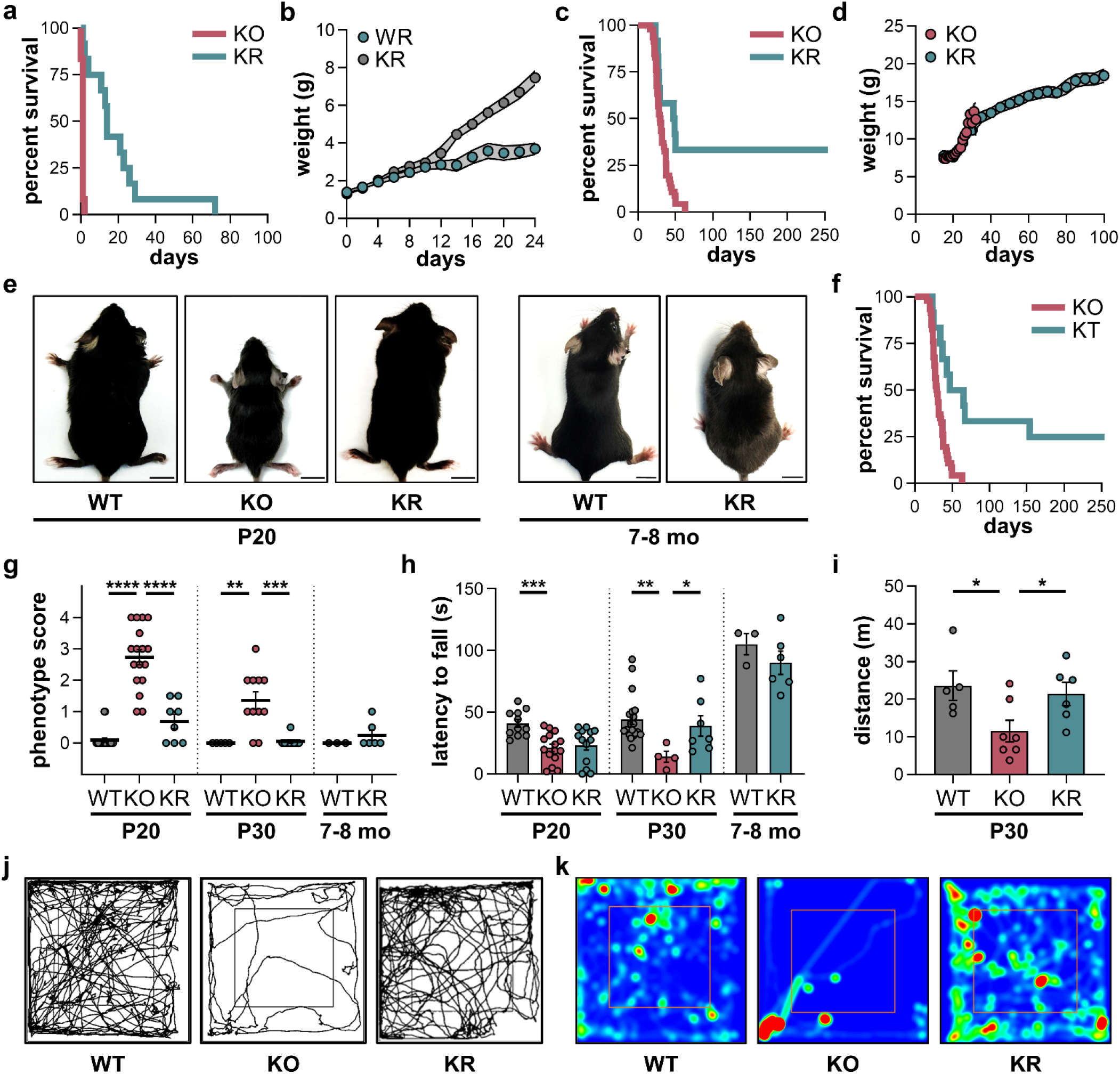
Rapamycin improves survival and disease phenotypes in MSUD mice. **a**, Survival curves of cMSUD mice treated with vehicle (KO, n=12, red) and rapamycin (KR, n=12, teal) starting at PND0 (4 mg/kg/day, i.p. injection). p<0.0001, log-rank. **b**, Weight progression of rapamycin treated wild type (WR, n=15, grey) and cMSUD (KR, n=15, teal) mice from PND0 to PND24. Grey shading represents ± SEM. **c, S**urvival curves of iMSUD mice treated with vehicle (KO, n=46, red) and rapamycin (KR, n=12, teal) starting at ∼PND15 (8 mg/kg/day, i.p. injection). p<0.0001, log-rank. **d**, Weight progression of iMSUD mice treated with vehicle (KO, n=46, red) or rapamycin (KR, n=12, teal) starting at PND15 (8 mg/kg/day, i.p. injection). Grey shading represents ± SEM. **e**, Representative images of WT, KO, and KR mice at PND20 (left) or at 7-8 months of age (right). Scale bar = 1 cm. **f**, Survival curves of iMSUD mice treated with vehicle (KO, n=46, red) or torin-2 (KT, n=12, teal). p<0.0001, log-rank. **g**, Phenotype scores (0-4) of WT, KO, or KR mice at PND20 (left), PND30 (middle), or at 7-8 months after treatment with vehicle or rapamycin. **h**, Latency to fall via rotarod assay of WT, KO, or KR mice at PND20 (left), PND30 (middle), or at 7-8 months after treatment with vehicle or rapamycin. **i**, Total distance traveled after 15-minutes in an open field test in WT, KO, or KR mice at PND30. Videos were analyzed using ANY-maze software. **j**, Representative open field track plots of WT, KO, or KR mice. Videos were analyzed using ANY-maze software. **k**, Representative heat maps of WT, KO, or KR mouse movement during open field test. Least to most time spent in a given area is represented by blue to red coloring, respectively. Unless noted otherwise, markers represent individual mice and results in charts are shown as mean ± SEM and statistical analysis was performed using a one-way ANOVA with Šídák multiple comparisons test. *p<0.05, **p<0.01, ***p<0.001, ****p<0.0001

Intermediate MSUD (iMSUD) mice were generated by Homanics by inserting a partially active liver-specific human BCKDH transgene into the cMSUD background^10^. This was achieved through transgenic insertion of the liver-specific tetracycline-controlled transactivator LAP promoter (LAP-tTa) and a human E2 variant of BCKDH with a tetracycline response element and alanine spacer (TRE-hBCKDH). As a result of partial BCKDH activity with the liver-specific hBCKDH transgene, these mice exhibit less severe BCAA and BCKA accumulation with a median lifespan of ∼30 days. This lifespan improves the practicality of working with a MSUD mouse model while allowing for robust phenotyping and assessment of therapeutic intervention. The cellular, biochemical, and anatomic phenotypes in iMSUD mice closely resemble human MSUD, arguably making iMSUD mice the most useful preclinical model to discover highly effective new therapies.

We treated iMSUD mice with vehicle or rapamycin. Rapamycin treatment increased survival with a maximal lifespan of >500 days compared to ∼30 days in control iMSUD mice whether initiated at PND15 or at weaning at PND21 (**Fig. 1c** and **Extended Data Fig. 1a**). Rapamycin-treated juvenile mice grew at slower rates than untreated WT mice, consistent with results in mitochondrial disease models^32^, however, weight gain continued to steadily increase until ∼6 months of age (**Fig. 1d** and **Extended Data Fig. 1b**). Rapamycin-treated mice were generally larger than vehicle-treated iMSUD mice and were largely indistinguishable from WT controls visually and in their behavior, especially in adulthood (**Fig. 1e**). Rapamycin is a potent inhibitor of mTOR Complex 1 (mTORC1) but it can sometimes indirectly inhibit mTOR complex 2^32^. Inhibition of mTORC2 may be harmful^36–38^, so we next asked whether direct mTOR inhibitors are similarly beneficial. An ATP-competitive inhibitor of mTORC1 and mTORC2 (torin-2) increased lifespan and weight similar to rapamycin (**Fig. 1f** and **Extended Data Fig. 1c**). Thus, direct inhibition of mTORC2 is not prohibitively harmful. It may alternatively have some beneficial effects as we previously reported in other mitochondrial disease mouse models^32^.

We modified a disease phenotype scoring system originally developed by Homanics (0=normal, 4=moribund) that assesses the presence or absence of common symptoms including body condition, muscle wasting, paralysis, tipping, wobbling, lateral recumbency, paw clasping, seizures, and other neurologic or muscular dysfunction observed in this model. The characteristic phenotype was visually apparent and severe in untreated iMSUD controls in cross-sectional studies nearing median lifespan (**Extended Data Fig. 1d** and **Video 1**), but not in rapamycin-treated mice. Noting this, we scored control and iMSUD mice treated with rapamycin (**Fig. 1g**). Consistent with literature, vehicle-treated iMSUD mice showed moderate to severe disease while WT mice appeared normal. Rapamycin treatment drastically improved the disease phenotype (**Fig. 1g**, **Extended Data Fig. 1d**, and **Video 1**), consistent with lifespan extension.

We next tested whether rapamycin improved neurobehavioral and motor coordination deficits in iMSUD mice. Control iMSUD mice showed time-progressive reductions in the latency to fall by the rotarod assay (**Fig. 1h**), a common readout for motor coordination and balance. Further, rapamycin restored rotarod performance that was sustained for >6 months (**Fig. 1h**). The iMSUD mice also showed a decline in locomotor activity by the open field test (**Fig. 1i-k** and **Supplementary Fig. S1**). Rapamycin treatment improved distance traveled, average moving speed, maximum speed, and exploratory behavior into central zones (**Fig. 1i-k**, **Extended Data Fig. 1e-g**). Control iMSUD mice had frequent lengthy immobile events and sporadic seizures which were not observed in rapamycin treated mice. Collectively, these data demonstrate rapamycin treatment greatly improves disease prognosis and neurologic behavior in iMSUD mice.

### Rapamycin reduces brain pathology and neuroinflammation

Noting the improvements in neurobehavioral performance, disease phenotypes, and survival in iMSUD mice upon rapamycin treatment, we next asked whether rapamycin similarly attenuates brain pathology. Cerebral edema is a life-threatening complication of MSUD and a common cause of death in MSUD patients during metabolic decompensation^17^. Clinical imaging demonstrates regional brain abnormalities involving the thalamus, basal ganglia, and myelination^39^. Previous studies in iMSUD mice identified neuropathology in the striatum and thalamus^40^. Conventional MRI did not reveal overt structural lesions (**Extended Data Fig. 2a** and **Supplementary Fig. S2**). To evaluate cerebral edema, we performed T2-weighted MRI in WT and iMSUD mice treated with vehicle or rapamycin. Vehicle-treated iMSUD mice showed increased age-progressive T2-weighted signal in the thalamus and similar trends in striatum (**Fig. 2a-b**, **Extended Data Fig. 2b-c**, and **Supplementary Fig. S3**), consistent with increased tissue water content and edema. The increase in regional T2-weighted signal was not observed in rapamycin-treated iMSUD mice (**Fig. 2a-b, Extended Data Fig. 2b-c**, and **Supplementary Fig. S3**).

**Figure 2.**
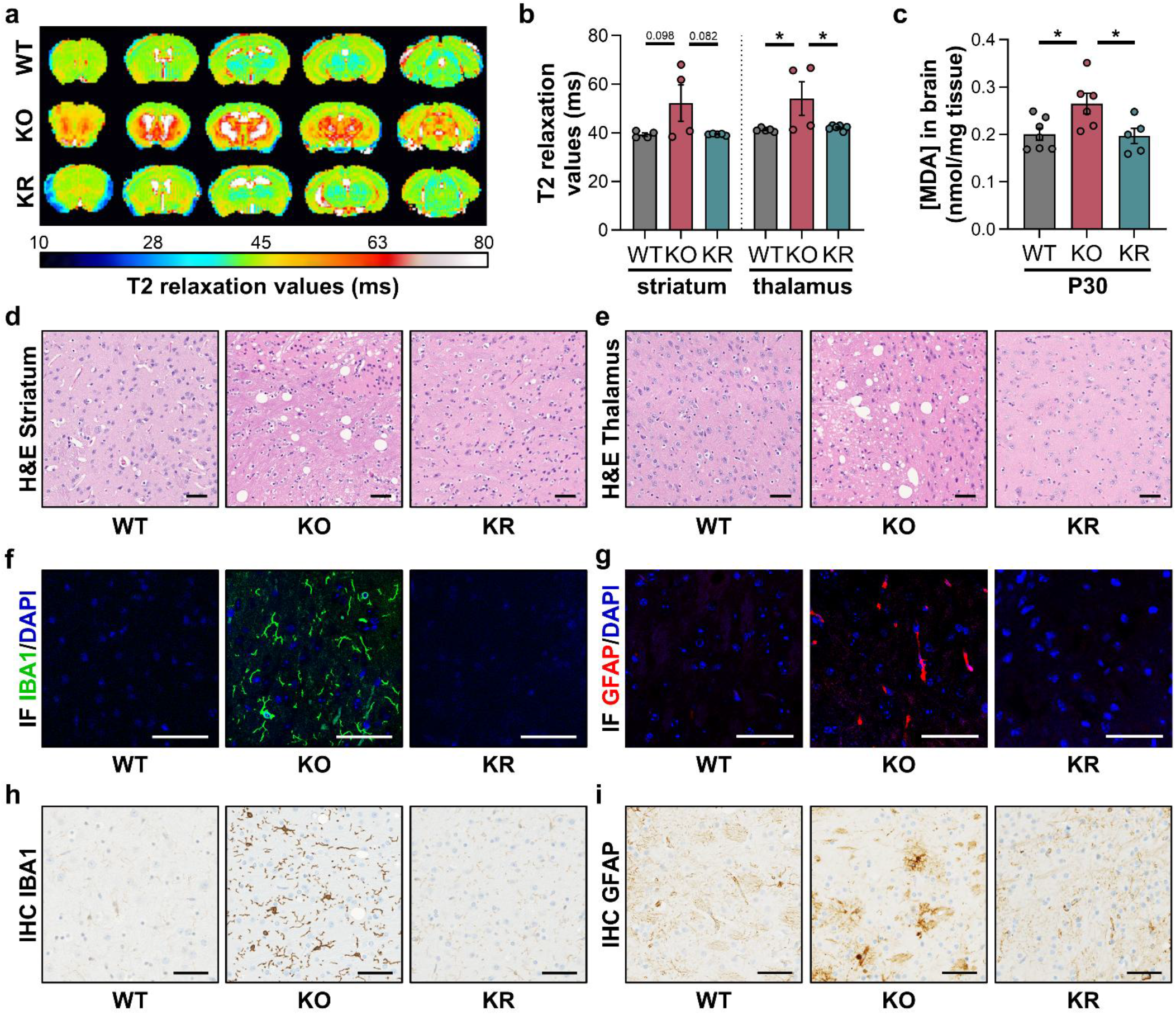
Rapamycin improves brain pathology and neuroinflammation in iMSUD mice. **a**, Representative MRI images of T2 relaxation times in WT, KO, or KR brain regions from rostral (left, olfactory bulb) to caudal (right, cerebellum) cross-sections at PND30. **b**, Quantified T2 relaxation values in striatum and thalamus of WT, KO, or KR mice at PND30. **c**, MDA concentration via a TBARS assay in whole brain isolated from WT, KO, or KR mice at PND30. **d, e**, Representative hematoxylin and eosin (H&E) histological stains assessing vacuolation and neuropathology in striatum or thalamus of WT, KO, or KR mice at PND30. Scale bars = 50 μm. **f, g**, Representative images of immunofluorescence (IF) of IBA1 (green), GFAP (red), and DAPI (blue) in WT, KO, or KR thalamus at PND30. Scale bar = 50 μm. **h, i**, Representative immunohistochemistry (IHC) of IBA1 and GFAP using DAB staining in WT, KO, or KR thalamus at PND30. Scale bar = 50 μm. Unless noted otherwise, markers represent individual mice and results in charts are shown as mean ± SEM and statistical analysis was performed using a one-way ANOVA with Šídák multiple comparisons test. *p<0.05

MSUD patient-derived cells and MSUD animals are reported to display markers of ROS-mediated toxicity including decreases in cellular glutathione pools and increased lipid peroxidation or DNA damage markers that are associated with oxidative stress and cell death^41^. Using whole brain tissue isolated from PND30 WT and iMSUD mice, we quantified levels of malondialdehyde using a MDA-TBARS assay as a readout for lipid peroxidation. We observed iMSUD mice have increased levels of MDA (**Fig. 2c**), consistent with reports in other MSUD models^41^. Rapamycin prevented this feature of oxidative damage in the brain (**Fig. 2c**). We next asked whether this was associated with brain pathology via histology (**Fig. 2d-e**, **Extended Data Fig. 3a-e**, and **Supplementary Fig. S4-9**). H&E staining revealed widespread vacuolation with highest frequency and size spread throughout the thalamus but was especially prominent on the periphery near the reticular nucleus adjacent to the internal capsule (**Fig. 2d-e**, **Extended Data Fig. 3a-b**, and **Supplementary Fig. S4**). Additionally, shrunken cells with intensely eosinophilic cytoplasm and condensed nuclei were observed in these regions, consistent with acute metabolic injury and neuronal degeneration. Dense regions of vacuolation also compromised the basal ganglia, specifically affecting the globus pallidus and to a lesser extent the caudate-putamen (**Fig. 2d**, **Extended Data Fig. 3d**, and **Supplementary Fig. S5-6**). The corpus callosum exhibited prominent rarefaction in iMSUD mice relative to WT controls (**Extended Data Fig. 3c-d** and **Supplementary Fig. S7**). Sporadic vacuoles were observed in the reticular formation (**Extended Data Fig. 3d** and **Supplementary Fig. S8**) while the hypothalamus, hippocampus, pons, and cerebral cortex were largely preserved at this age. Rapamycin greatly reduced spongiform pathology and neuronal injury in the thalamus and other affected brain regions (**Fig. 2d-e**, **Extended Data Fig. 3a-d**, and **Supplementary Fig. S4-9**). This neuroprotective effect largely persisted for >6 months of continuous rapamycin treatment, though spongiform pathology in the medulla was apparent in some mice (**Extended Data Fig. 3e** and **Supplementary Fig. S9**).

Our observations of cerebral edema by T2-weighted MRI, elevated lipid peroxidation, and histopathological abnormalities collectively suggested ongoing tissue injury in the iMSUD brain. Given the well-established interplay between oxidative stress, cellular damage, and neuroimmune activation, we hypothesized that these pathological changes would be associated with neuroinflammation. Previous reports in iMSUD mice showed astrocytosis^40^, consistent with abnormal dendritic morphology in MSUD patients^42^. This led us to examine cellular markers of glial activation to determine whether inflammatory processes contribute to MSUD neuropathology and whether rapamycin treatment modifies these responses. We quantified glial fibrillary acidic protein (GFAP) and ionized calcium-binding adapter molecule 1 (IBA1), which are well-established markers of astrocyte and microglial activation, respectively. In untreated iMSUD mice whole brain extracts, both GFAP and IBA1 protein levels were increased by WB relative to WT controls (**Extended Data Fig. 4a-b**), reflecting enhanced reactive astrogliosis and microglial activation consistent with neuroinflammation. In contrast, rapamycin treatment led to a significant reduction in both GFAP and IBA1, suggesting that rapamycin suppresses neuroinflammation (**Extended Data Fig. 4a-b**).

Comprehensive immunohistochemical analysis was performed to assess whether increased GFAP and IBA1 expression was consistent with increased frequency of glia or if it was also associated with morphological changes consistent with inflammation. We first performed immunofluorescence with multiple coronal sections to evaluate cell number (**Fig. 2f-g**, **Extended Data Fig. 4c-d**, and **Supplementary Fig. S10-11**). We observed widespread increases in GFAP- and IBA1-positive signal in most brain regions. Consistent with our previous results, we observed the largest increases in the number of GFAP-positive and IBA1-positive cells in the thalamus, and to a lesser extent the basal ganglia, in iMSUD mice, but not with rapamycin treatment (**Fig. 2f-g**, **Extended Data Fig. 4c-d**, and **Supplementary Fig. S10-11**). Immunohistochemistry was used for morphological analysis, in which we once again observed an increase in the number of glial cells (**Fig. 2h-i**, **Extended Data Fig. 4e-f**, and **Supplementary Fig. S12-13**). Beyond their increased abundance, astrocytes and microglia in iMSUD controls exhibited morphological alterations consistent with neuroinflammation. Within the thalamus, GFAP-positive astrocytes showed hypertrophic morphologies with thickened, star-shaped processes, while IBA1-positive microglia displayed enlarged cell bodies and we observed an increased frequency of amoeboid morphologies suggesting a pro-inflammatory state (**Fig. 2h-i** and **Supplementary Fig. S12-13**). These reactive morphological features were prominently reduced following rapamycin treatment and were sustained for >6 months (**Fig. 2h-i** and **Extended Data Fig. 4g**). Collectively, these data strongly support rapamycin improves brain pathology, physiology, and function in iMSUD mice.

### Rapamycin mitigates skeletal muscle pathology

Skeletal muscle dysfunction is an increasingly recognized clinical manifestation of MSUD in patients^11–16^. Similar to patients, iMSUD mice experience skeletal muscle dysfunction including hypotonia, myofiber necrosis, myofibrillar damage, and decreased muscular performance^11–16^. We therefore asked whether rapamycin improves muscle physiology. Functional assessment using grip strength and inverted grid hanging tests revealed impaired performance in iMSUD mice relative to WT controls at early disease stages (**Fig. 3a-b**). Although rapamycin only slightly improved performance at these early points, a robust improvement became evident with age. At older ages, rapamycin-treated iMSUD mice exhibited grip strength and hanging performance comparable to WT (**Fig. 3a-b**).

**Figure 3.**
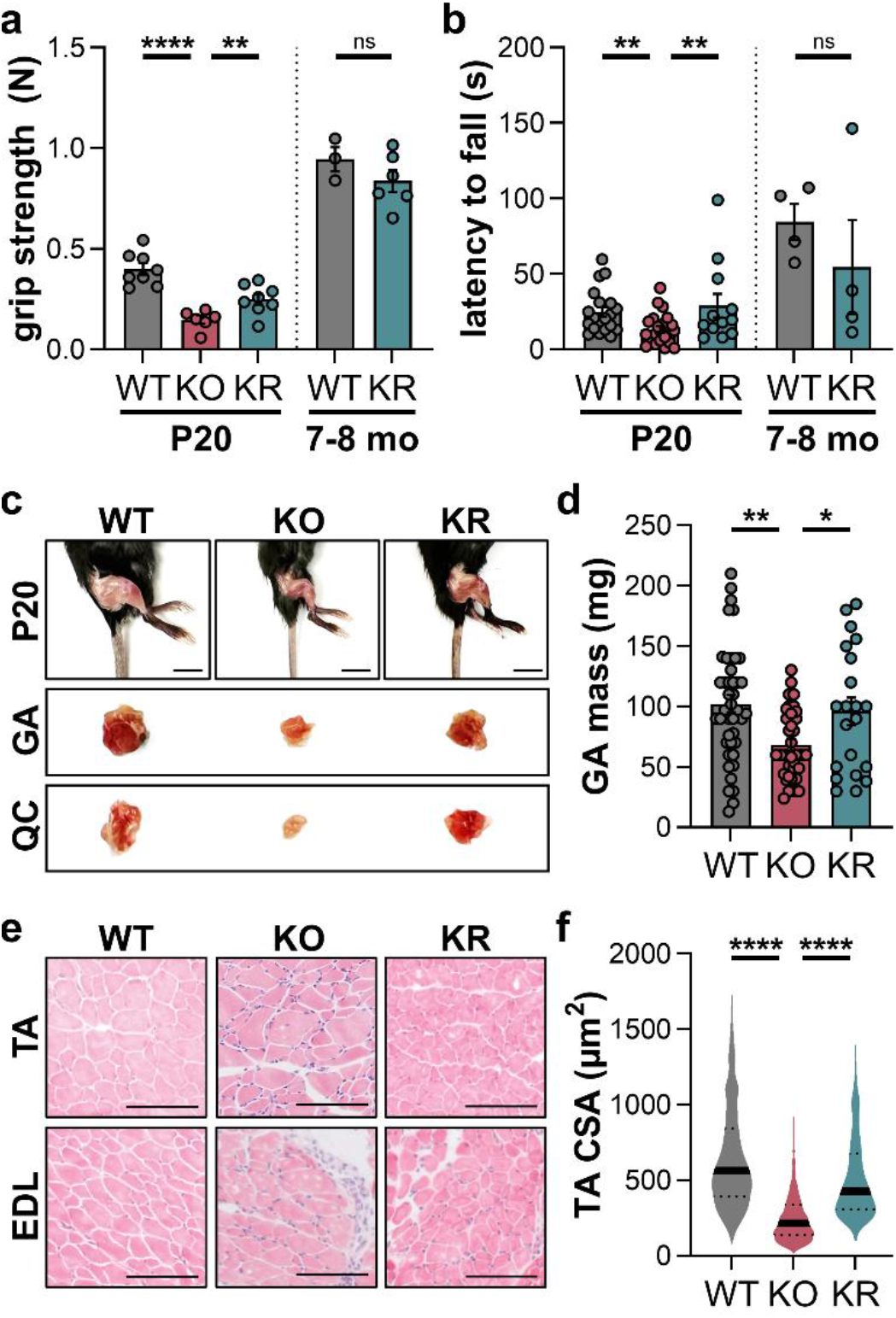
Rapamycin rescues muscle pathology. **a**, Peak forelimb grip strength measured across indicated ages (PND20 and 7-8 month) in WT, KO, or KR mice. **b**, Inverted grid hanging (wire hanging) test measured across indicated ages listed previously. **c**, Representative hind limb skeletal muscle images and isolated muscle images of quadriceps (QC) and gastrocnemius (GA) from WT, KO, or KR mice. Scale bar = 1 cm. **d**, Weight of gastrocnemius muscles isolated from WT, KO, or KR mice at PND20. **e**, Representative H&E staining of tibialis anterior (TA) and extensor digitorum longus (EDL) muscle in WT, KO, or KR mice at PND20. Scale bar = 100 μm. **f**, Analysis of TA myofiber cross-sectional area in WT, KO, or KR mice at PND20. Statistical analysis of CSA (n=246-2453 fibers counted) was performed using a non-parametric Kruskal-Wallis test with Dunn’s multiple comparisons test. Unless noted otherwise, markers represent individual mice and results in charts are shown as mean ± SEM and statistical analysis was performed using a one-way ANOVA with Šídák multiple comparisons test. p<0.01, ****p<0.0001, ns=not significant

Consistent with this, rapamycin increased muscle size and mass (**Fig. 3c-d**). H&E staining of tibialis anterior (TA) and extensor digitorum longus (EDL) muscles revealed hallmarks of pathology in iMSUD mice but not with rapamycin (**Fig. 3e**). Cross-sectional area analysis of myofibers in TA and EDL revealed a significant decrease in myofiber size distribution in iMSUD mice compared to WT (**Fig. 3f** and **Extended Data Fig. 5**). The iMSUD mice exhibited visible loss of muscle and fat mass at these ages reaching moribundity, coinciding with hindlimb paralysis and consistent with muscle wasting, atrophy, or cachexia. This decrease was partially rescued in iMSUD mice treated with rapamycin. For some vehicle-treated iMSUD mice, TA and EDL hypotonia was so severe that cross-sectional areas could not be reliably obtained. Together, these results demonstrate that rapamycin produces durable neurologic and neuromuscular rescue, not merely lifespan extension.

### Rapamycin lowers BCAAs and BCKAs without restoring BCKDH activity

We believe a central challenge to the development of effective medications to treat MSUD is a limited understanding of BCAA pathobiology. Our results with rapamycin suggest mTOR is involved, however, other pathways have been implicated. To begin to test these mechanisms, we assessed survival of iMSUD mice when screened with interventions that target these proposed pathways. It has been proposed that deficiencies in mitochondrial bioenergetics through disrupted TCA cycle anaplerosis and reductions in brain glutamate levels may cause neuron toxicity from high leucine^40^. We used the α-ketoglutarate precursor, dimethylketoglutarate (DMKG), which has been shown to improve cellular energetics and increase brain α-ketoglutarate levels supporting normal glutamate/glutamine metabolism^43^. We used doses of DMKG used in other mouse models of mitochondrial disease^43^, yet it had no effect on median or maximal lifespan (**Extended Data Fig. 6a**). Tryptophan deficiency is one of the longest standing mechanistic hypotheses proposed in neurotoxicity in MSUD due to inhibition of tryptophan uptake into neurons via LAT1 and disruptions in serotonin signaling^44^. Consistent with this, norleucine blocks leucine uptake in the brain and showed some efficacy in MSUD mouse models^40^. The brain permeable prodrug α-methyl-tryptophan is used as a PET tracer for studying serotonin synthesis and can functionally boost serotonin signaling^44^. However, treatment of iMSUD mice with this precursor had no effect under our tested conditions (**Extended Data Fig. 6b**). In contrast to the use of mTOR inhibitors, 17-AAG (protein degradation), hypoxia (mitochondrial dysfunction), and PI3K inhibition (insulin signaling) also had no positive effects under our tested conditions (**Extended Data Fig. 6c-e**).

We thus investigated the capacity for rapamycin to inhibit mTOR in iMSUD mice. As mTOR is a nutrient-responsive kinase that alters phosphorylation of proteins involved in cell growth, metabolism, and in other factors, we focused on evaluating protein expression of mTOR targets and phospho-proteins by immunoblot at earlier stages of disease before median lifespan of iMSUD mice is reached (PND20). Phosphorylation of S6 ribosomal protein and AKT are well-established markers of mTORC1 and mTORC2 activity, respectively. Using brain protein extracts, we observed that iMSUD controls showed slight increases in mTORC1 activation using phosphorylated S6 as a robust marker for mTORC1 (**Fig. 4a-b**), but did not observe drastic changes in mTORC2. Consistent with the role of rapamycin as a potent mTOR inhibitor, we were unable to observe phospho-S6 expression in brain of rapamycin-treated mice (**Fig. 4a-b**).

**Figure 4.**
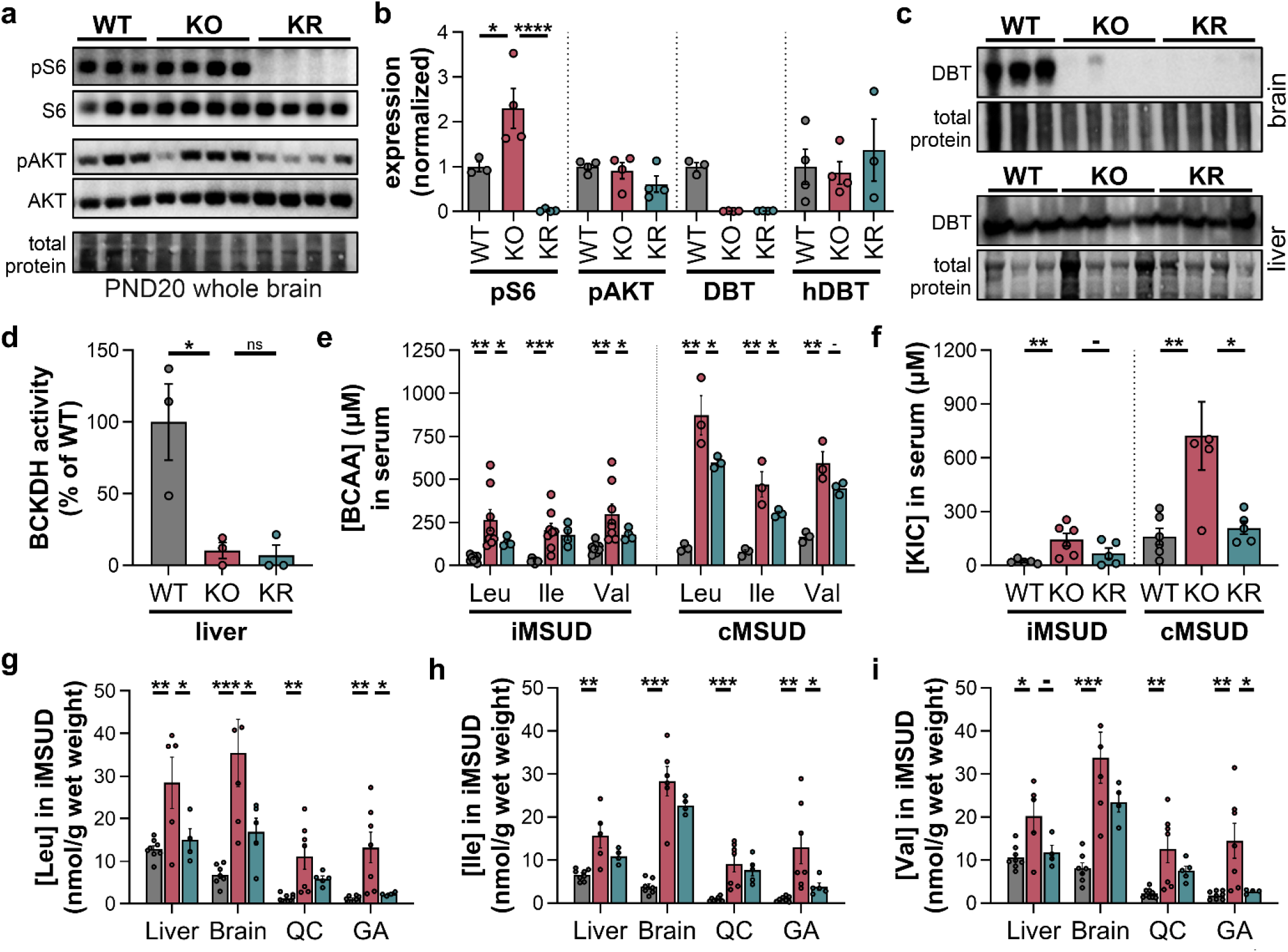
Rapamycin reduces BCAA accumulation independent of BCKDH. **a**, Representative western blot (WB) images of pS235-S6, S6, pS473-AKT, and AKT in whole brain extracts from WT, KO, or KR mice at PND20. **b**, Densitometry of WBs relative loading control. **c**, WB images of DBT in iMSUD brain (top) and liver (bottom) isolated from WT, KO, or KR mice at PND20. Statistical analysis was performed using a one-way ANOVA with Šídák multiple comparisons test. **d**, Relative BCKDH activity in liver tissue from WT, KO, or KR mice at PND30, shown as % of WT activity. **e**, Absolute concentrations of the BCAAs leucine, isoleucine and valine in WT (grey) and iMSUD (PND20, red) or cMSUD KO (PND1, red) or KR (teal) sera by HPLC after AQC derivatization. **f**, Absolute concentrations of the BCKA ketoisocaproate (KIC) in WT, iMSUD, or cMSUD sera by HPLC after OPD derivatization. **g-i**, Quantification of absolute levels of leucine, isoleucine, or valine in WT (grey), KO (red), or KR (teal) iMSUD mice at PND20 in liver, brain, quadriceps (QC), and gastrocnemius (GA) normalized to tissue weight. Unless noted otherwise, markers represent individual mice and results in charts are shown as mean ± SEM and statistical analysis was performed using a one-tailed student’s t-test with Bonferroni correction based on literature precedent for increased BCAAs in MSUD models. **-**p<0.10, *p<0.05, **p<0.01, ***p<0.001, ****p<0.0001, ns=not significant

As we described previously, iMSUD mice were generated by Homanics by selectively expressing a modified and partially active human DBT transgene in the liver of cMSUD mice with a global deletion of murine DBT (the E2 subunit of BCKDH). To evaluate whether rapamycin was effective in iMSUD mice by altering BCKDH, we quantified DBT expression and BCKDH activity. We observed that rapamycin had no effect on DBT protein expression in liver of iMSUD mice, and unsurprisingly DBT was still missing in the brain (**Fig. 4b-c**). Reversible phosphorylation of BCKDHA, the E1 subunit of BCKDH, at Ser293 by BCKDH kinase (BCKDK) inhibits BCKDH activity. BCKDK helps tune BCKDH activity in response to BCAA levels through mechanisms involving tyrosine kinases, insulin signaling, allosteric regulation from BCKA status, and ATP/ADP ratio^45,46^. We evaluated BCKDHA phosphorylation as mTOR could indirectly modulate its phosphorylation. Using liver tissue extracts, we observed a decrease in relative BCKDHA phosphorylation in iMSUD normalized to WT controls (**Extended Data Fig. 6f-g**). Rapamycin treatment increased pS293-BCKDHA phosphorylation, which would alternatively suggest rapamycin treatment reduces BCKDH activity. Rapamycin also had no meaningful effects on the expression of branched-chain amino transferases (BCAT1 and BCAT2) that reversibly convert BCAAs to their corresponding BCKAs in brain extracts (**Extended Data Fig. 6f-g**). Further, we quantified relative activity of BCKDH using a biochemical assay in liver isolated from iMSUD mice. Consistent with previous reports^10^, we observed a ∼90% reduction in liver BCKDH activity in iMSUD controls relative to WT liver extracts (**Fig. 4d**). Rapamycin did not improve BCKDH activity (**Fig. 4d**). These results led us to conclude that the improvements in iMSUD prognosis with rapamycin are independent of BCKDH activity.

The observed changes in phosphorylation of BCKDHA are consistent with positive feedback resulting from high serum BCAA in control iMSUD mice that is reduced upon rapamycin treatment. As all pathogenic mechanisms converge on high BCAA and BCKA status, we finally quantified BCAAs and BCKAs in iMSUD mice at PND20. Absolute BCAA levels in serum, brain, liver, and skeletal muscle were quantified using HPLC after AQC derivatization and UV-Vis detection and the BCKA ketoisocaproate (KIC) was quantified via similar methods after derivatization with *o*-phenylenediamine (OPD). We observed a global accumulation of leucine, isoleucine, and valine as well as their corresponding ketoacids in iMSUD mice in all tissues tested (**Fig. 4e-i** and **Extended Data Fig. 6h**). Strikingly, rapamycin treatment led to general reductions in the BCAAs and KIC (**Fig. 4e-i** and **Extended Data Fig. 6h**). Heat maps generated using the z-scores obtained from the absolute quantification data clearly demonstrated rapamycin treatment strongly reduced BCAA and KIC status (**Extended Data Fig. 6i**). Even a single dose of rapamycin was able to reduce BCAA and KIC levels within 24 hours in serum of cMSUD mice at PND1 (**Fig. 4e-f**). Thus, rapamycin rapidly reduces the toxic accumulation of BCAAs.

## DISCUSSION

We discovered that rapamycin extends survival and attenuates disease progression in leading mouse models of classic and intermediate MSUD. We observed that rapamycin reduced brain pathology, glial reactivity, and skeletal muscle degeneration. Rapamycin lowered BCAAs and BCKAs without restoring DBT expression or hepatic BCKDH activity. Strikingly, even a single dose of rapamycin decreased circulating BCAAs and BCKAs in cMSUD neonates. Previous studies suggest that BCAA accumulation disrupts brain amino acid balance and neurotransmitter availability^18,40^. Excess BCKAs impair mitochondrial bioenergetics^19^. More recently, AAV and mRNA approaches demonstrated that directly restoring BCKDH is effective in treating MSUD mice^35,47,48^. Our results represent a complementary strategy that has similar efficacy profiles. However, the liver-restricted transgenic expression of BCKDH underlying the iMSUD model differs from MSUD patients, and efficacy in MSUD mice does not establish responsiveness in genetically diverse human populations. It is critical to determine whether rapamycin is effective in more human-relevant systems. However, the biochemical reduction of BCAA and BCKA status with rapamycin provides a compelling rationale for testing its broader applicability.

The broad protection of brain structure and function further suggests that rapamycin is acting on an important mechanism of MSUD neuropathology. Previous studies in iMSUD mice identified cerebral edema, vacuolation, altered brain amino acid and neurotransmitter levels, and pathology in the thalamus and striatum^40^. Clinical imaging similarly shows that MSUD encephalopathy affects the thalamus, globus pallidus, internal capsule, brainstem, and other myelinated regions, with diffusion abnormalities in these areas commonly attributed to intramyelinic edema^39^. Consistent with these findings, we observed increased T2 signal, widespread vacuolation in the thalamus, pathology in the globus pallidus, caudate-putamen, and reticular formation, rarefaction of the corpus callosum, oxidative damage, and reactive astrogliosis and microglial activation. This pattern is consistent with a regionally selective metabolic encephalopathy in iMSUD mice with prominent vacuolation in the thalamus, increased gliosis in the thalamus and basal ganglia, and additional abnormalities involving myelinated axons or other white matter processes. Disruptions in the basal ganglia and thalamus provide a strong anatomical basis for the progressive wobbling, tipping, hindlimb dysfunction, and impaired motor coordination we observed in iMSUD mice. It further suggests perturbations in GABAergic or glutamatergic signaling underlie leucine toxicity, consistent with depletions in GABA and glutamate levels in MSUD^18,40^. Rapamycin prevented these abnormalities. We do not yet know whether rapamycin acts directly in neurons or whether brain protection occurs primarily through the periphery. However, the simultaneous improvement in brain structure, glial reactivity, neurologic function, and systemic metabolism suggests that rapamycin is modifying the underlying disease process rather than only alleviating one consequence of MSUD.

The robust response to rapamycin suggests that mTOR is not simply providing an off-ramp from an isolated feature of MSUD pathology. Rather, mTOR signaling may be integrated into disease pathophysiology. Rapamycin-treated iMSUD mice were largely indistinguishable from WT controls across survival, disease phenotype, neurologic function, and tissue pathology, indicating a remarkably broad improvement in disease progression. The mechanism of leucine toxicity in modulating mTOR activity remains unclear, and our studies with interventions for other proposed mechanisms do not definitively rule them out. We observed an increase in phosphorylation of the mTORC1 target S6 in iMSUD brain, possibly identifying mTOR hyperactivation directly because of leucine sensor regulation. However, it is possible that the intracellular localization of mTOR or its binding profiles, not its direct enzymatic activity, may also be perturbed in MSUD. It remains unknown which downstream mTOR effectors mediate the rescue, particularly because mTOR regulates diverse processes including protein synthesis, autophagy, lipid metabolism, inflammation, and cellular stress responses. These mechanisms may differ between the brain and skeletal muscle, where the requisite mTOR activity coupled with the metabolic demands and pathological responses to BCAA accumulation are distinct. We believe the rescue reflects a fundamental metabolic adaptation, consistent with the broad effects of rapamycin on amino acid and lipid metabolism previously observed in other mitochondrial disease models.

An alternative framework underlying our findings is that MSUD pathology may arise not solely from the metabolic toxicity of leucine, but also from maladaptive nutrient sensing. The field has proposed leucine accumulation as a cytotoxic metabolic feature of MSUD that disrupts neurotransmitter balance, mitochondrial bioenergetics, redox homeostasis, and other cellular processes. While these mechanisms are supported by strong experimental evidence, they do not preclude nutrient stress signaling as an important contributor to disease. Leucine is among the most potent physiological regulators of mTOR signaling^27–29^, raising the possibility that leucine not only serves as a toxic metabolite in MSUD, but also as an aberrant nutrient signal that persistently alters cellular responses to amino acid availability. Under this model, pathology results from both the direct biochemical consequences of BCAA accumulation and unintended activation of nutrient-responsive pathways that normally sense fluctuations in amino acid status. Consistent with this, a recent report in *Bckdk*-knockout mice suggests reducing leucine via fasting suppresses mTORC1 misregulation^49^. The strong rescue with rapamycin is consistent with interruption of these maladaptive signaling responses being sufficient to attenuate disease progression. More broadly, these findings suggest that nutrient-sensing pathways may contribute to the pathophysiology of other inherited metabolic disorders characterized by chronic metabolite accumulation, consistent with the efficacy of rapamycin in multiple disorders associated with mitochondrial dysfunction^32^.

The in-depth mechanism by which rapamycin lowers BCAAs also remains unknown. Our findings argue against increased DBT abundance or restoration of BCKDH activity, but do not distinguish altered intestinal absorption, tissue utilization, or excretion of BCAAs. Reduced intestinal absorption from the diet could decrease exogenous BCAA delivery. Altered protein synthesis or protein degradation could change the availability of endogenous BCAAs. Rapamycin treatment could also alter BCAA incorporation into protein, residual metabolic flux, or excretion of BCAAs and BCKAs from the body. BCAAs can serve as terminal electron acceptors^50^ and previous work showed that metformin lowers KIC in patient-derived fibroblasts and iMSUD mice alongside reduced muscle BCAT expression^30^, demonstrating that metabolite production upstream of BCKDH is itself pharmacologically modifiable. Rapamycin did not alter BCAT abundance in our experiments, suggesting a distinct mechanism. Unraveling this mechanism requires in-depth and rigorous studies, individually probing each of these pathways.

Rapamycin possesses a practical advantage as a repurposed drug that may streamline translation. Sirolimus (also known as rapamycin), sold under the brand name Rapamune, has been FDA-approved for decades, is available orally, and has extensively characterized pharmacodynamics, pharmacokinetics, drug interactions, and monitoring requirements. There exists an established infrastructure for sourcing and manufacturing, clinical distribution, and quality control standards. These advantages could lower many barriers to clinical development, but approval for other diseases does not establish safety or efficacy in MSUD. Low dose sirolimus was tested clinically in slow flow vascular malformations, including in 33 children^51^. From this study, there was a reported clinical benefit at target concentrations of low dose sirolimus (4–10 ng/mL) and fewer serious adverse events were reported than with higher doses (10-15 ng/mL). Rapamycin is being investigated in late-stage clinical trials in children with tuberous sclerosis complex. A low dose (1 mg tablets) is reported to reduce epileptic seizure frequency without harming patient growth and development^52^. Our observation of slower MSUD growth with rapamycin demonstrates the need to identify the lowest exposure to rapamycin that remains efficacious with less unwanted effects on growth or development. Patients with mTOR hyperactivity may alternatively be less susceptible to rapamycin-mediated toxicity and side effects.

An important next step is to evaluate rapamycin alongside current best management practices. Current management strategies for MSUD combine dietary leucine restriction with frequent biochemical outpatient monitoring. These approaches improve nutritional control of leucine status, but do not fully prevent neurological and psychiatric complications. Similarly, liver transplantation restores hepatic BCKDH activity but is unable to reverse brain damage that has already occurred, and patients sometimes require repeated liver transplantations. Rapamycin should be tested for additive benefits under controlled nutritional conditions, including whether it improves metabolic stability without compromising amino acid availability. Prevention of metabolic crisis is a particularly important question because catabolic illness can drive encephalopathy despite careful dietary leucine management.

More broadly, our findings suggest that nutrient-responsive signaling is an important mediator in the pathophysiology of MSUD. However, this work also extends beyond MSUD. BCAA-derived metabolites can influence organ transport of lipids and influence insulin resistance^8^, and BCAA restriction is increasingly recognized as critical in the effects of caloric restriction in normative aging and longevity. Thus, the study of MSUD provides a powerful system to define how nutrient signaling controls pathological amino acid burden, with the potential to uncover therapeutic principles relevant to other inherited metabolic diseases and acquired disorders of amino acid homeostasis.

## METHODS

### Animals and Animal Use

Animal procedures were approved by the Institutional Animal Care and Use Committee (IACUC) at the University of Cincinnati (protocol 25-07-31-01) and were conducted in full compliance with the guidelines outlined by the National Institutes of Health (NIH) for the Care and Use of Laboratory Animals. Mice were housed in a temperature-controlled environment within a University of Cincinnati animal facility on a 14:10 hr light/dark cycle. Heterozygous breeder pairs were generously donated by the Kennedy Lab (Buck Institute) to establish an iMSUD mouse colony. cMSUD mouse breeders could not be obtained externally and were instead generated by backcrossing iMSUD heterozygous mice with wild type C57Bl/6J mice to remove the transgenes as reported recently^35^. Pups produced by pairwise breeding of *Dbt^+/-^* mice were genotyped by PCR before post-natal day 21 (PND21) to produce *Dbt^+/+^*, *Dbt^+/-^*, or *Dbt^-/-^* pups. Male and female mice were distributed evenly in experiments, and iMSUD or cMSUD mice were housed with at least one healthy littermate of the same sex for companionship and warmth. A maximum of five mice were housed in each cage. Mice were monitored daily by laboratory staff for disease progression, weight loss, and moribundity. Routine animal husbandry, including daily monitoring for health status, food and water intake, and daily cage/food changes were carried out by trained animal facility staff. Humane endpoints were strictly observed, and animals exhibiting a ≥20% loss in body weight or signs of severe distress, low body condition, or moribundity were euthanized using IACUC-approved methods.

### Drug treatment

iMSUD mice received daily intraperitoneal (i.p.) injections beginning at PND15-21, unless otherwise indicated, and cMSUD mice beginning at PND0. Drugs were administered at 6.66 µL per gram body weight, with injection sides alternated daily. Compounds were generally dissolved in DMSO, sequentially mixed with PEG-400 and Tween-80, and diluted in 0.9% saline or PBS. Rapamycin (A2B Chem, AB46980) was administered to iMSUD mice at 8 mg/kg (1.2 mg/mL solution) and cMSUD mice at 4 mg/kg (0.6 mg/mL solution). Treatment initiation at PND15, or PND21 produced similar outcomes, but all functional, histological, and biochemical studies in this work began at PND15. Torin-2 (Chemenu, CM106094) was administered at 1 mg/kg (0.15 mg/mL solution) from PND15-20 and 5 mg/kg (0.75 mg/mL solution) thereafter in 3% DMSO, 10% PEG-400, and 5% Tween-80. Additional treatments included α-methyltryptophan (Santa Cruz Biotechnology, sc-300182, 25 mg/kg from PND21, 1% DMSO/PBS), 17-AAG (MedChemExpress, HY-10211, 10 mg/kg from PND17, 1% DMSO/10% PEG-400/10% Tween-80), dimethylketoglutarate (MedChemExpress, HY-44134, 300 mg/kg from PND10, PBS), and dactolisib (MedChemExpress, HY-50673, 5 mg/kg from PND15, 3% DMSO/40% PEG-400/5% Tween-80). Vehicle contained 1% DMSO, 5% PEG-400, and 5% Tween-80 in saline. Solutions were passed through a 0.22-µm sterile filter and stored frozen before use. For hypoxia treatment, iMSUD mice were maintained continuously at 11% O₂ starting from PND25 in custom acrylic chambers supplied by a HigherPeak MAG-20 generator.

### Phenotype Scoring

Periodically, mice were evaluated for their phenotype score representing illness severity in MSUD. This was achieved by a five-point phenotype scoring system based on disease symptom severity, with a score of 0 representing normal and unencumbered movements, 1 representing minor phenotypes of gait disturbance, tipping and motor deficiency that do not preclude the mouse’s ability to access food and water, 2 representing greater severity of gait disturbance, seizures, and muscle wasting resulting in difficulty accessing food and water readily, 3 representing life-threatening hindlimb paralysis, lateral recumbency, labored breathing and failure to thrive, and 4 representing a moribund state which requires immediate humane euthanasia.

### Open Field Test

Mouse locomotor activity and exploratory behavior were analyzed by an open field test. Open-field testing was performed in a custom-built 20”x20” arena consisting of an HDPE base and opaque acrylic walls supplied from TAP Plastics. Mice were placed in the center of the arena and allowed to explore undisturbed and without noise for 15 minutes. Video was simultaneously captured from a tripod above. Videos were then analyzed using ANY-maze software for track plots, total distance traveled, average speed, etc.

### Rotarod Test

Motor coordination and balance in mice were assessed using a digital rotarod apparatus (IITC Life Science I-755 rotarod) with Series 8 software. Mice were placed on a horizontal rotating rod, and the speed gradually increased from 0 to 45 rpm. The rotarod test length was set to a maximum of 180 seconds. The latency to fall was recorded in seconds as a measure of motor coordination. Mice were trained 1-2 days before on how to use the rotarod, and three trials per mouse were used to determine average latency to fall for each individual mouse.

### Grip Strength Test

Assessment of muscle forelimb grip strength was performed using an automatic grip strength meter (Soonkoda digital force gauge S-3N). The mouse was allowed to grasp the grid and was gently pulled backward by the tail until it lost its grip. The peak force was captured by the force meter in newtons. An average of four trials of peak force were taken per mouse separated by ∼15 minutes each.

### Inverted Grid Hanging Test

Mice were placed on a ∼12”x12” stainless steel grid (6.0 mm openings, 0.7 mm thickness) and allowed to establish a secure grip before the grid was inverted, positioning the mice upside down approximately 18 inches above a soft surface. Latency to fall from the inverted grid was recorded as a measure of neuromuscular performance. Mice were trained 1-2 days before and at least three trials per mouse were used to determine average latency to fall for each individual mouse.

### Tissue Collection and Storage of Tissues

Mice were fed *ad libitum* and euthanized by unanesthetized cervical dislocation at the indicated age (PND20, PND30, or around 7-8 months of age). Immediately after euthanasia, blood and various tissues were quickly collected, snap-frozen in liquid nitrogen, and stored temporarily at low temperatures. For further processing, the frozen tissues were finely ground under liquid nitrogen using a chilled mortar and pestle. The powdered tissue was then transferred into pre-cooled cryogenic vials and stored in liquid nitrogen for long-term preservation and future experimental use. For tissues used for histology, mice were euthanized by CO2 asphyxiation after MRI imaging and tissues were carefully dissected, fixed in formalin, and stored in 70% ethanol or PBS before embedding by the Integrative Pathology Core Facility at Cincinnati Children’s Hospital Medical Center.

### H&E Staining

To evaluate neuropathological changes, iMSUD brain tissues were subjected to hematoxylin and eosin (H&E) staining at the CCHMC pathology core facility. Hematoxylin was used to stain the brain to visualize cell nuclei (purple color), and eosin was used to visualize the cytoplasm (pink color). To examine the presence of histopathological abnormalities like vacuolation, we examined the brain slides under a Leica DMi8 Widefield Fluorescence microscope and a Zeiss Axioscan Slide Scanner (for full coronal brain images and regional images). The images were analyzed using Fiji (ImageJ).

### Immunofluorescence

GFAP and IBA1 were evaluated in coronal brain slices of iMSUD brains in thalamus, hypothalamus, striatum, hippocampus, cortex, midbrain, cerebellum, and pons sections. PBS was used for incubations or perfusion overnight at 4°C. Brain tissues were paraffin-embedded, sectioned, and processed at the CCHMC pathology core facility. Primary antibodies like IBA1 (1:100, Cell Signaling Technology, 17198T,) and GFAP (Roche, 760-4345) were used and nuclei counterstained with DAPI, followed by an Alexa Fluor 488-conjugated goat anti-rabbit IgG (1:100, Invitrogen, A11008) and Alexa Fluor 594-conjugated goat anti-rabbit IgG (1:100, Invitrogen, A11037) secondary antibodies. Immunofluorescence images were captured via a confocal microscope using Microscope Leica Stellaris 8 Confocal and ImageJ was used for signal quantification. Images were analyzed using ImageJ on deidentified slides.

### Immunohistochemistry

Immunohistochemical analysis was performed on iMSUD brain tissues to assess neuroinflammation. Brain processing, paraffin embedding, and immunostaining were performed at the CCHMC pathology core facility. using IBA1 (1:100, Cell Signaling Technology, 17198T) and GFAP (Roche, 760-4345) and detection was performed using the ready-to-use Discovery UltraMap DAB anti-Rabbit detection kit (Roche, 05266564001), containing the Discovery ChromoMap DAB kit (Roche, 05266645001). Histopathological images were captured using a Leica DMi8 Widefield Fluorescence microscope and Zeiss Axioscan Slide Scanner (for full coronal brain images and regional images) and analyzed using ImageJ software on deidentified slides.

### T2-Weighted MRI Imaging

MRI mapping of iMSUD mice was performed at PND20, PND30, and in 7-8-month-old mice using a vertical wide-bore 9.4 T Bruker Avance III HD MRI equipped with a gradient insert and a 36 mm proton volume coil. Mice were anesthetized using 2% isoflurane in 2 L/min oxygen during the scan. Each mouse was scanned using axial Fast Spin Echo sequences for T2-weighted (TR = 4000 ms, effective TE = 71.5 ms, RARE factor = 20, echo space= 6.5 ms, 4 averages) images. Fat saturation and motion averaging were applied. Fifteen interlaced slices were imaged with the following parameters like slice thickness 0.75 mm, 0.3 mm gap, FOV=2.84×2.84 cm with 192×192 matrix. Based on the T2 images, 5 slices were centered over the potential location for T2 mapping using a multi-slice multi-echo sequence (TR = 2000 ms, 20 echoes from TE = 9.21 ms to 184.14 ms at 9.207 ms echo space, 2 averages). Images were acquired with the following parameters: slice thickness 0.75 mm, 0.75 mm gap, FOV = 2.84×2.84 cm with 142×142 matrix. A Bo map was used to improve reconstruction, and parametric images of T2 relaxation were produced using the Bruker Paravision software. Further analysis was conducted using Imalytics preclinical software. The 15-slice T2W images were aligned with the mouse brain Beneveniste Mirrione MRI atlas, and then the 5-slice T2 signal and map images were aligned to the atlas using the same transform. A mask was then created to strip and segment the T2 map data. Average T2 relaxation values were recorded for brain regions of interest (e.g. thalamus and striatum). For presentation, the stripped T2 map was displayed using the Royal color LUT set from 10 ms–80 ms and saved in TIFF as a montage above the corresponding 5-slice T2 signal images.

### Protein Extraction

Protein was extracted from pulverized tissues in RIPA buffer (Thermo Fisher Scientific 89901) containing protease/phosphatase inhibitors (Sigma-Aldrich PPC1010) at roughly 20 µL/mg tissue. Homogenization was performed using a mechanical homogenizer or rapid sonication followed by incubation at 4°C for one hour with repeated re-homogenizations. After incubation, samples were centrifuged at 17,000×g for 15 minutes at 4°C to separate the supernatant from the tissue pellet. The resulting supernatants were collected and used for protein quantification and analysis. Protein extracts were stored at -80°C before use.

### Immunoblot

Protein concentrations were determined by a BCA assay according to manufacturer instructions. Samples were adjusted to 1-2 mg/mL in RIPA buffer, mixed with LDS sample buffer (GenScript M00676) and 4% β-mercaptoethanol (Bio Basic, MB0338) as reducing agent. Samples were denatured at 95°C for 5 minutes. Proteins (30 μg per well) were separated on 4-12% Bolt^TM^ Bis-Tris Plus gels using MOPS running buffer (Thermo Fisher B000102) and transferred to PVDF membranes using the Trans-Blot Turbo Transfer System (Bio-Rad) according to manufacturer instructions with the addition of 0.2% SDS to the transfer buffer. A molecular weight marker (Thermo Fisher Scientific 26616) was used as ladder. Total protein was visualized using No-Stain^TM^ Protein Labeling Reagent (Thermo Fisher A44449) according to manufacturer instructions. After rinsing, membranes were blocked with 5% BSA in TBST for 30 min, incubated with primary antibodies (at the dilutions listed below) in 5% bovine serum albumin (BSA, Fisher Scientific BP1600) overnight at 4°C followed by incubation at room temperature for 1-2 hrs. Membranes were repeatedly washed with TBST (3×10 mins) and incubated with HRP-conjugated secondary antibody (at the dilutions listed below) in 5% BSA at room temperature for 2 hrs. Membranes were repeatedly washed with TBST and chemiluminescence was detected using the SuperSignal^TM^ Pico PLUS (Thermo Fisher Scientific 34578) according to manufacturer instructions with an iBright 1500 imaging system. Protein expression was analyzed by densitometry using ImageJ relative to total protein or actin and normalized to WT controls. Primary antibodies included phospho-S6 ribosomal protein (Ser235/236) (1:1,000, Cell Signaling Technology, 4858S), S6 ribosomal protein (1:1,000, Cell Signaling Technology, 2217T), phospho-AKT (Ser473) (1:4,000, Cell Signaling Technology, 4060S), total AKT (1:4,000, Cell Signaling Technology, 4691S), GFAP (1:1,000, Cell Signaling Technology, 12389S), IBA1 (1:1,000, Cell Signaling Technology, 17198T), DBT (1:2,500, GeneTex, GTX113412), and HRP-conjugated β-actin (1:20,000, Cell Signaling Technology, 5125S). The corresponding secondary antibody for each primary included donkey anti-rabbit HRP-conjugated secondary antibody (1:20,000 in 5% BSA; Thermo Scientific 31458), HRP-conjugated mouse antibody binding protein (1:2,000) in 5% BSA; Santa Cruz sc-516102), or horse anti-mouse HRP-conjugated secondary antibody (1:20,000 in 5% BSA; Cell Signaling cs-7076).

### BCAA Quantification

Branched-chain amino acids were quantified in serum, brain, liver, quadriceps, and gastrocnemius muscle by HPLC following derivatization with an excess of 6-aminoquinolyl-N-hydroxysuccinimidyl carbamate (AQC). Approximately 20 mg of frozen powdered tissue or 10 µL serum was extracted in methanol/water (4:1, v/v) following homogenization in water/PBS (9:1, v/v). Samples were vortexed, incubated at -20°C for 30 min, sonicated cold for 30 min, and centrifuged (17,000 x g, 15 min, 4°C). Supernatants were collected, dried under vacuum with a SpeedVac, reconstituted in borate buffer (100 mM boric acid at pH=8.8, ∼1.5 µL/mg tissue used) and derivatized by adding freshly prepared AQC reagent (∼60 µg used per reaction from freshly prepared 10 mM stock in dry acetonitrile). Samples were vortexed followed by 10-fold dilution with borate buffer and centrifugation. Supernatant was collected and filtered via syringe filter into HPLC vials. Samples were analyzed by HPLC with UV detection at 254 nm using an Eclipse Plus C18 column (USDBB03390, Agilent B15064, 2.1×150 mm, RRHD 1.8 µm) with water/0.1% formic acid and acetonitrile/0.1% formic acid eluents. Metabolites were separated with a step-gradient from 100% water to 85% water over 12 minutes, followed by a continuous gradient to 65% water after 28 minutes, before switching to 100% acetonitrile for 3 minutes. Leucine, isoleucine, and valine concentrations were determined by comparison to external calibration curves generated from authentic amino acid standards (0.01-100 µM) processed in parallel with experimental samples.

### BCKA Quantification

Branched-chain keto acids were quantified in serum, brain, liver, and skeletal muscle by HPLC following derivatization with *o*-phenylenediamine (OPD). Tissue samples (∼20 mg) were homogenized and extracted using methanol/water and water/PBS as described previously, while serum (10 µL) was processed directly. Following centrifugation and solvent removal by SpeedVac, samples were derivatized with OPD by reconstituting in 20% acetic acid (90 µL) followed by addition of OPD (10 µL of a 10 mM stock). Samples were vortexed and gently heated at 90°C for 30 mins. Derivatized BCKAs were extracted into ethyl acetate after diluting with water, evaporated to dryness, reconstituted in DMSO, filtered via syringe filter, and analyzed by HPLC at 350 nm using an Eclipse Plus C18 column (USDBB03390, Agilent B15064, 2.1×150 mm, RRHD 1.8 µm) with water/0.1% formic acid and acetonitrile/0.1% formic acid eluents. Metabolites were separated with a step-gradient from 100% water to 85% water over 2 minutes, followed by a continuous gradient to 30% water after 10 minutes, before switching to a continuous gradient back up to 100% water after 25 minutes. Quantification of α-ketoisocaproate (KIC), α-keto-β-methylvalerate (KMV), and α-ketoisovalerate (KIV) was performed using external calibration curves generated from authentic standards processed in parallel with experimental samples. Absolute quantification of KMV and KIV were not reliable due to low limits of detection, and thus the absolute concentration data were not included due to low confidence in the accuracy using this analytical method. However, relative changes were robust and thus the data was used for z-score analysis.

### MDA-TBARS Assay

Lipid peroxidation was evaluated by measuring malondialdehyde (MDA) concentrations in mouse brain tissue using a TBARS assay kit (Cayman Chemical 10009055). Brain tissue (∼25 mg) was homogenized in 250 µL RIPA buffer containing protease inhibitors and centrifuged at 1,600 × *g* for 10 min at 4 °C. The resulting supernatant was collected for analysis. An MDA standard curve ranging from 0 to 50 µM was prepared from a 125 µM stock solution using the supplied MDA standard (Cayman Chemical 10009202). For each sample or standard, 100 µL was combined with 100 µL SDS solution and 800 µL color reagent containing thiobarbituric acid, acetic acid, and sodium hydroxide according to manufacturer instructions. Reactions were incubated at 95 °C for 1 hr, immediately cooled on ice for 5 min and centrifuged at 1,600 × *g* for 10 min at room temperature. The resulting supernatant (200 µL) was transferred to a 96-well plate, and absorbance was measured at 535 nm. MDA concentrations were calculated from the standard curve.

### BCKDH Activity Assay

Fresh liver tissue (∼50 mg) was homogenized in 10 µL/mg tissue weight in cold extraction buffer using a MagnaLyser at 6,000 rpm. Homogenates were centrifuged at 20,000 × g for 5 min at 4°C, and proteins were precipitated with 9% PEG for 20 min on ice. Following centrifugation at 12,000 × g for 10 min at 4°C, pellets were resuspended in 5 µL/mg tissue weight suspending buffer. Samples were added to prewarmed assay buffer, and absorbance at 340 nm was recorded for 40 min at 37°C to determine background activity. α-Ketoisovalerate was then added to a final concentration of 1 mM, and NADH formation was monitored for an additional 40 min. Background-corrected reaction slopes were normalized to initial tissue weight and expressed relative to WT activity. Buffers were prepared as previously described^53^.

### Muscle Cross-Sectional Area

Mouse muscle histology was analyzed similar to previously described^54^. TA and EDL muscles were dissected as part of whole mouse lower limbs. After degloving, limbs were placed in Tissue-Tek O.C.T. and frozen by floating tissue molds on supercooled 2-methylbutane over liquid nitrogen and stored at - 80°C. Cross-sections of murine lower limbs were collected on a Leica CM1850 cryostat at -20°C. Muscle was sectioned at 10 μm thickness and collected every 100 μm along the length of the limb. Sections were then stained with H&E to identify general morphology and quantify myofiber cross-sectional area (CSA). Sections were imaged with a 10X phase contrast objective lens on an Olympus IX83 inverted microscope. Similar areas on each skeletal muscle were chosen for imaging, using the tibia and fibula as landmarks. Myofiber CSA was measured in μm^2^ using ImageJ on deidentified slides. Between 200-1500 myofibers were analyzed per biological replicate.

### Statistical Analysis

All data are presented as means ± SEM. Pathologic analysis of brain and muscle were performed on deidentified slides. Statistical analyses and plots were made using GraphPad Prism software (version 10.6.1, GraphPad Software, San Diego, CA, USA). Statistical significance was assessed using Student’s t-test with appropriate correction for multiple comparisons (e.g. Bonferroni correction), one-way ANOVA followed by Šídák’s multiple-comparisons test, Kruskal-Wallis test followed by Dunn’s multiple-comparisons test, or log-rank test, as indicated in the figure legends. **-**p<0.10; *p < 0.05; **p < 0.01; ***p < 0.001; ****p < 0.0001; ns = not significant

## DATA AVAILABILITY

All raw source data is provided with this paper.

## CODE AVAILABILITY

No custom code was used in this work.

## ACKNOWLEDGEMENTS

We thank Alessandro Bitto, Yuehmei Hsu, Matt Kaeberlein, Brian Kennedy, and Arvind Ramanathan for helpful discussions, guidance, and assistance with preliminary work on MSUD. We thank Sena Asefa, Camille Bodart, Reid Couch, Alyssa Del Castillo, Vivian Ha, Sydney Huff, Divine Idahosa, CJ Kelly, Megan McGuire, Paige Miller, Ruth Negash, Karlee Parry, Josh Poole, Ellie Puente, Ava Scherz, Isaiah Smith, Sushila Thapa, and other undergraduate students for assistance with genotyping mice, experimental support, and/or in daily mouse handling. We thank Temugin Berta and Artur Prudente for assistance and guidance with rotarod experiments. We thank Fatoumata Dabo for assistance with running the HPLC for BCAA quantification through the ACS SEED program. We thank Eric Treinen for assistance with months of frequent rapamycin treatment. We thank Lisa Lemen and Sharon Wang (Preclinical Imaging Core, College of Medicine, UC) for the MRI mapping acquisition. We thank Chet Closson (Live Microscopy Core, CoM, UC) for his technical assistance with microscopy instrumentation used for H&E, IHC, and IF imaging. We thank Betsy DiPasquale and Drew Smith (Pathology Research Core, CCHMC) for their assistance with brain tissue processing, paraffin embedding, and preparation of histological (H&E) and immunostaining (IHC and IF) slides.

## AUTHOR CONTRIBUTIONS

ASG conceived the study, designed experiments, performed experiments, analyzed and interpreted data, provided funding, and wrote and edited the paper. SG designed and performed most experiments, analyzed and interpreted data in most experiments, and wrote and edited the paper. MES designed, performed, and analyzed experiments on BCAA/BCKA quantification and immunoblot, analyzed muscle histology, and edited the paper. IAA performed experiments and analyzed data on mouse behavioral studies and cMSUD lifespan. APH designed, performed, and analyzed experiments for BCKDH activity assay and immunoblot, and edited the paper. RKD performed procedures in mice and in tissue collections. KC analyzed data for muscle histology. KEV designed and performed experiments and analyzed or interpreted data for muscle histology. SG, MES, IAA, APH, and RKD performed most day-to-day mouse handling, observations, breeding, data collection in live mice, and performed tissue collections.

## SUPPLEMENTARY INFORMATION

**Supplemental Figure 1.** Open field test track plots of multiple mice replicates in WT, KO, and KR groups at PND20.

**Supplemental Figure 2.** MRI images of multiple mice replicates in WT, KO, and KR groups at PND20, PND30, and 7-8 months.

**Supplemental Figure 3.** T2 mapping of multiple mice replicates in WT, KO, and KR groups at PND20, PND30, and 7-8 months.

**Supplemental Figure 4.** Expanded view of thalamus of multiple mice replicates in WT, KO, and KR groups at PND30 and 7-8 months.

**Supplemental Figure 5.** Expanded view of striatum of multiple mice replicates in WT, KO, and KR groups at PND30 and 7-8 months.

**Supplemental Figure 6.** Expanded view of the globus pallidus of multiple mice replicates in WT, KO, and KR groups at PND30 and 7-8 months.

**Supplemental Figure 7.** Expanded view of the corpus callosum of multiple mice replicates in WT, KO, and KR groups at PND30 and 7-8 months.

**Supplemental Figure 8.** Expanded view of the midbrain of multiple mice replicates in WT, KO, and KR groups at PND30 and 7-8 months.

**Supplemental Figure 9.** Expanded view of the pons/medulla of multiple mice replicates in WT, KO, and KR groups at 7-8 months.

**Supplemental Figure 10.** Images of multiple mice replicates of immunofluorescence staining for GFAP in WT, KO, and KR groups at PND30 and 7-8 months.

**Supplemental Figure 11.** Images of multiple mice replicates of immunofluorescence staining for IBA1 in WT, KO, and KR groups at PND30 and 7-8 months.

**Supplemental Figure 12.** Images of multiple mice replicates of immunohistochemical staining for GFAP in WT, KO, and KR groups at PND30 and 7-8 months.

**Supplemental Figure 13.** Images of multiple mice replicates of immunohistochemical staining for IBA1 in WT, KO, and KR groups at PND30 and 7-8 months.

**Video 1**. Representative video of neurological symptoms present (tipping/tremors) in iMSUD KO mice (middle panel, phenotype score=2.5) compared to WT (left panel, phenotype score=0) and KR (right panel, phenotype score=0.5).

## SOURCE DATA

All raw source data, including uncropped/unedited western blot images, has been provided with this manuscript in supplemental files.

## EXTENDED DATA FIGURES

**Extended Data Figure 1.**
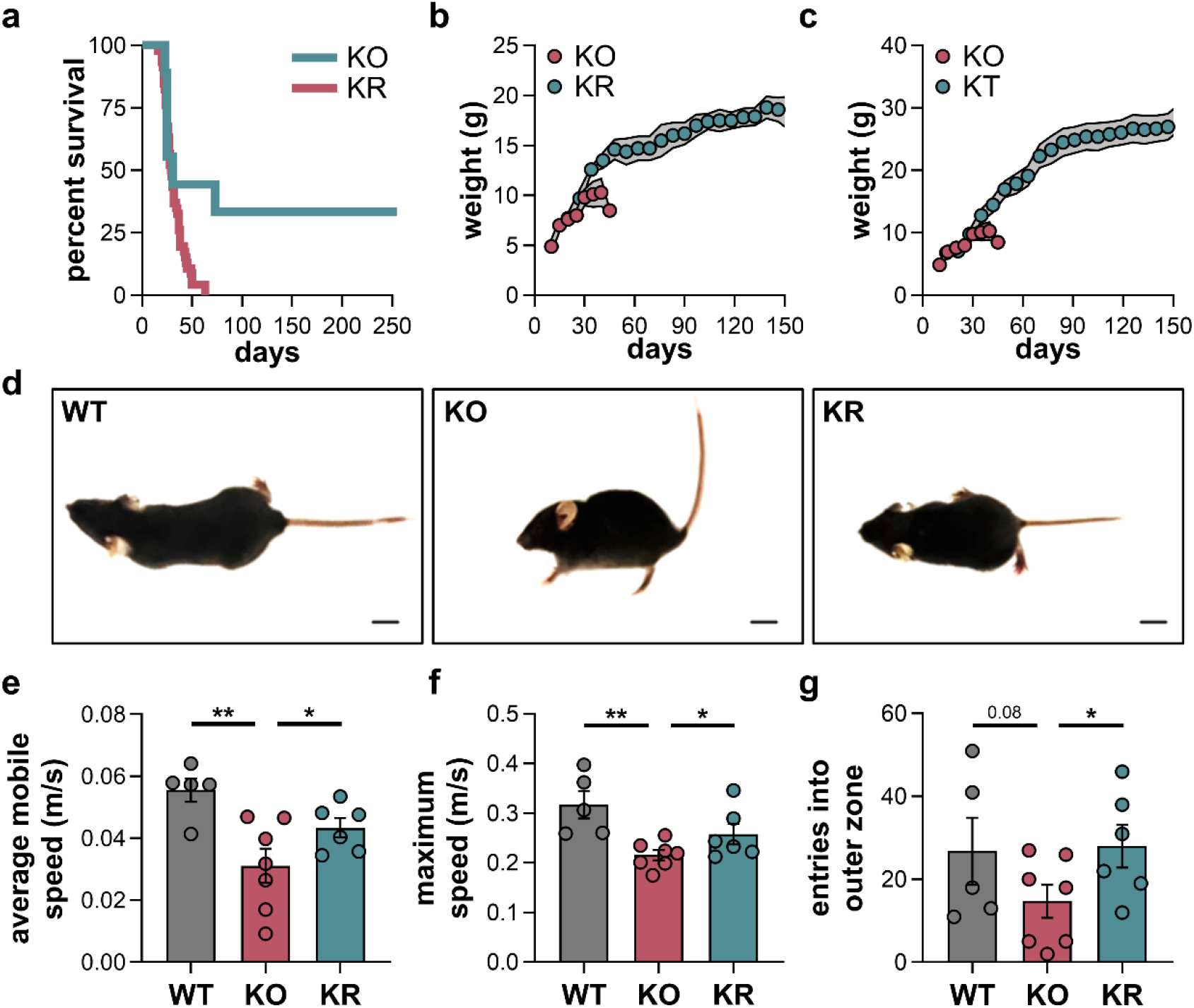
Rapamycin improves survival and disease phenotypes in iMSUD mice. **a**, Survival curves of iMSUD mice treated with vehicle (KO, n=46, red) and rapamycin (KR, n=9, teal) starting at PND21 (8 mg/kg/day, i.p. injection). p<0.0001, log-rank. **b**, Weight progression of iMSUD mice treated with vehicle (KO, n=46, red) or rapamycin (KR, n=9, teal) starting at PND21. Grey shading represents ± SEM. **c**, Weight progression of iMSUD mice treated with vehicle (KO, n=46, red) or torin-2 (KT, n=12, teal) starting at PND15. Grey shading represents ± SEM. **d**, Representative images of phenotypic illness associated with WT (left), KO (middle), and KR (right) iMSUD mice at PND30. Scale bar = 1 cm. **e**, average mobile speed in a 15-minute open field test of WT, KO, and KR iMSUD mice at PND30. Videos were analyzed using ANY-maze software. **f**, maximum speed in a 15-minute open field test of WT, KO, and KR mice. Videos were analyzed using ANY-maze software. **g**, outer zone entries made in a 15-minute open field test of WT, KO, and KR iMSUD mice. Unless noted otherwise, markers represent individual mice and results in charts are shown as mean ± SEM and statistical analysis was performed using a one-way ANOVA with Šídák multiple comparisons test. *p<0.05, **p<0.01

**Extended Data Figure 2.**
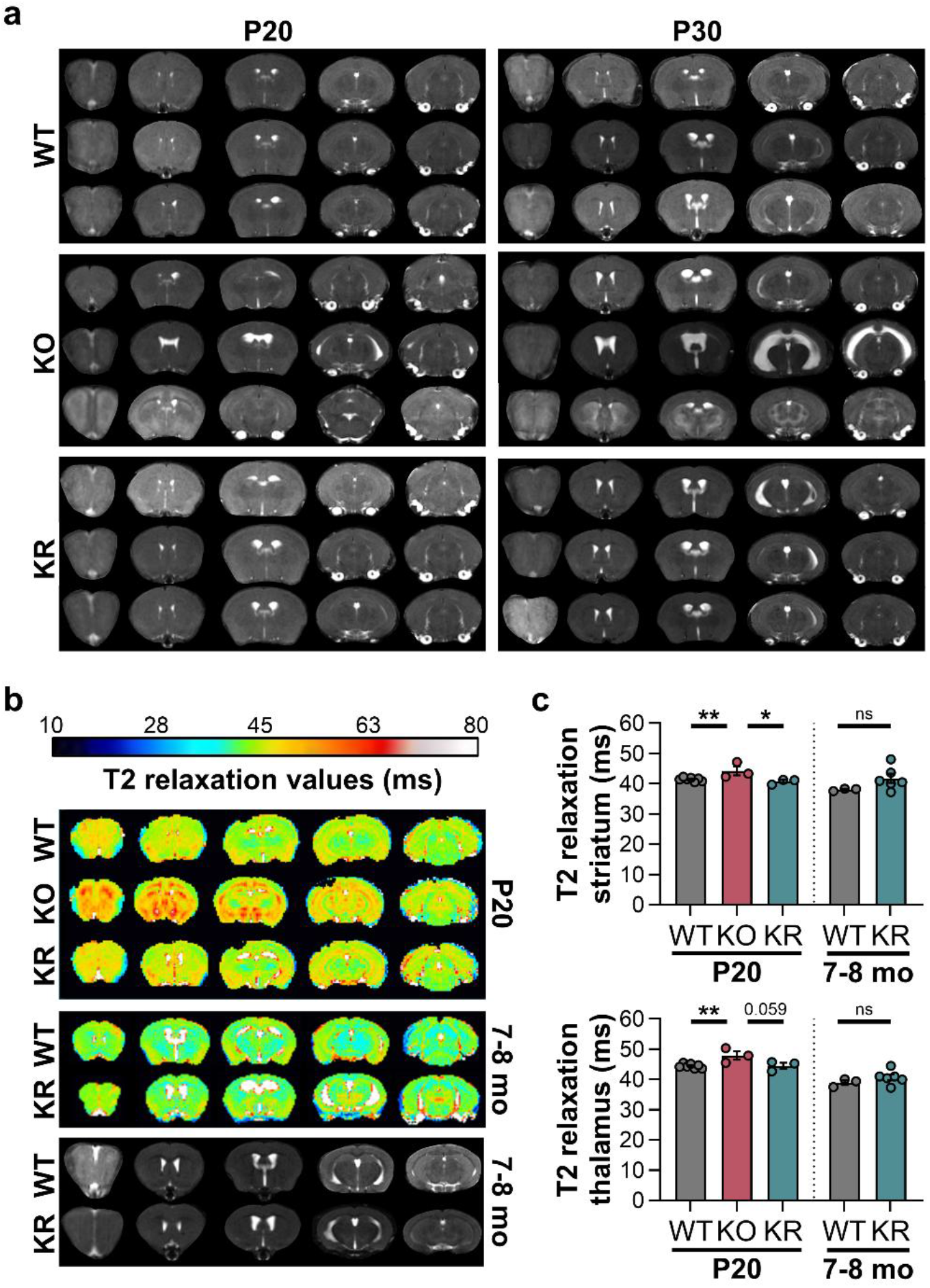
Rapamycin prevents cerebral edema. **a**, Representative MRI images in WT, KO, or KR brain regions from rostral (left, olfactory bulb) to caudal (right, cerebellum) cross-sections at PND20 and PND30. **b**, Representative MRI images of T2 relaxation times in WT, KO, or KR brain regions from rostral (left, olfactory bulb) to caudal (right, cerebellum) cross-sections at PND20 and 7-8 months. **c**, Quantified T2 relaxation values in striatum and thalamus of WT, KO, or KR mice at PND20 and 7-8 months. Unless noted otherwise, markers represent individual mice and results in charts are shown as mean ± SEM and statistical analysis was performed using a one-way ANOVA with Šídák multiple comparisons test. *p<0.05, **p<0.01

**Extended Data Figure 3.**
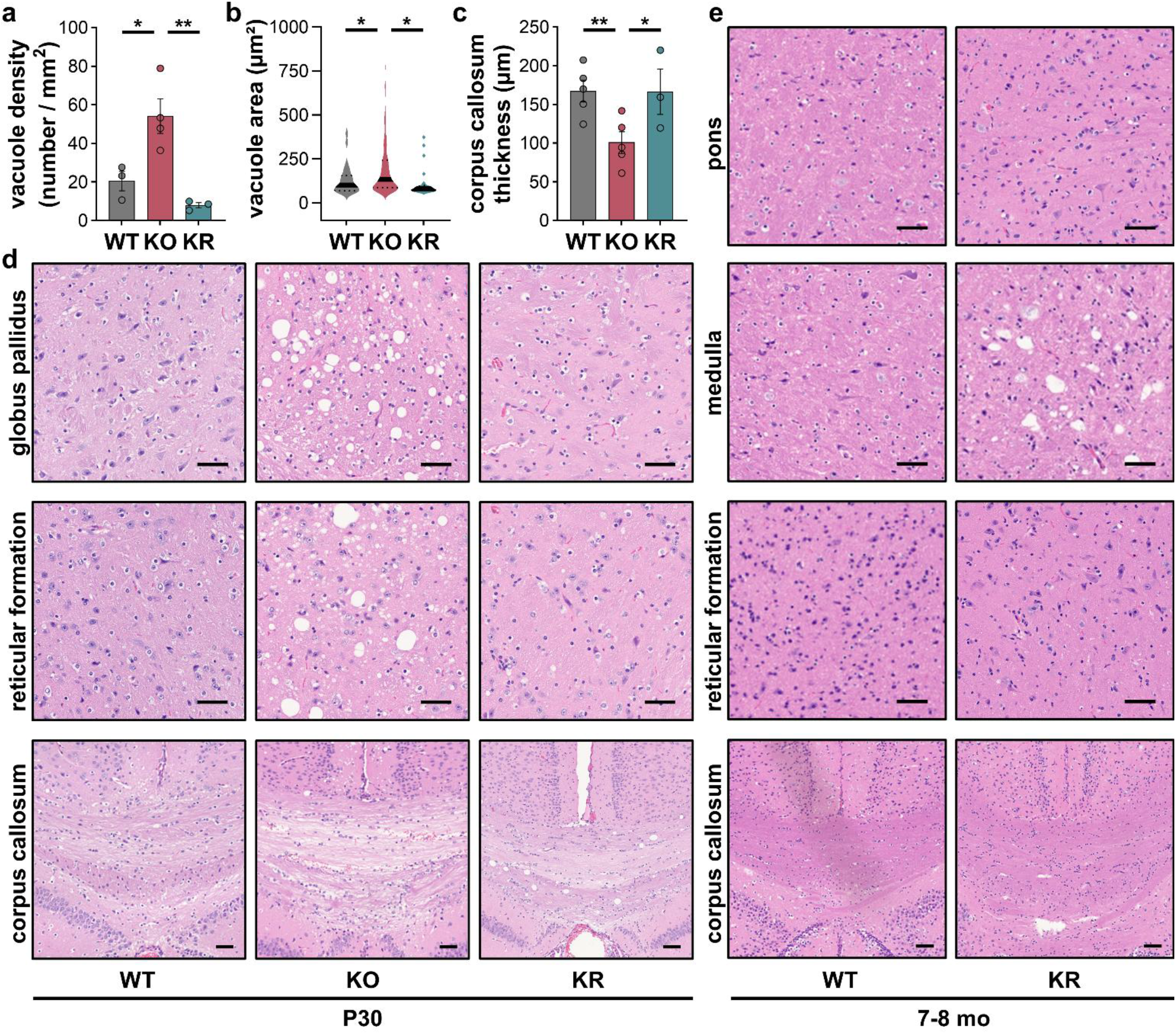
Rapamycin rescues brain pathology. **a**, Quantification of vacuolation frequency in H&E staining of the thalamus in WT, KO, and KR iMSUD brain. **b**, Quantification of vacuole size in H&E staining of the thalamus in WT (n=26 vacuoles from 3 mice), KO (n=343 vacuoles from 3 mice), and KR (n=22 vacuoles from 3 mice) iMSUD brain. **c**, Quantification of average thickness of the midline of the midbody of the corpus callosum in H&E staining of WT, KO, and KR iMSUD brain. **d**, Representative H&E staining of the globus pallidus, reticular formation, and corpus callosum in WT, KO, or KR mice at PND30. Scale bar = 50 μm. **e**, Representative H&E staining of the pons, medulla, reticular formation, and corpus callosum in WT or KR mice at 7-8 months. Scale bar = 50 μm. Unless noted otherwise, markers represent individual mice and results in charts are shown as mean ± SEM and statistical analysis was performed using a one-way ANOVA with Šídák multiple comparisons test. *p<0.05, **p<0.01

**Extended Data Figure 4.**
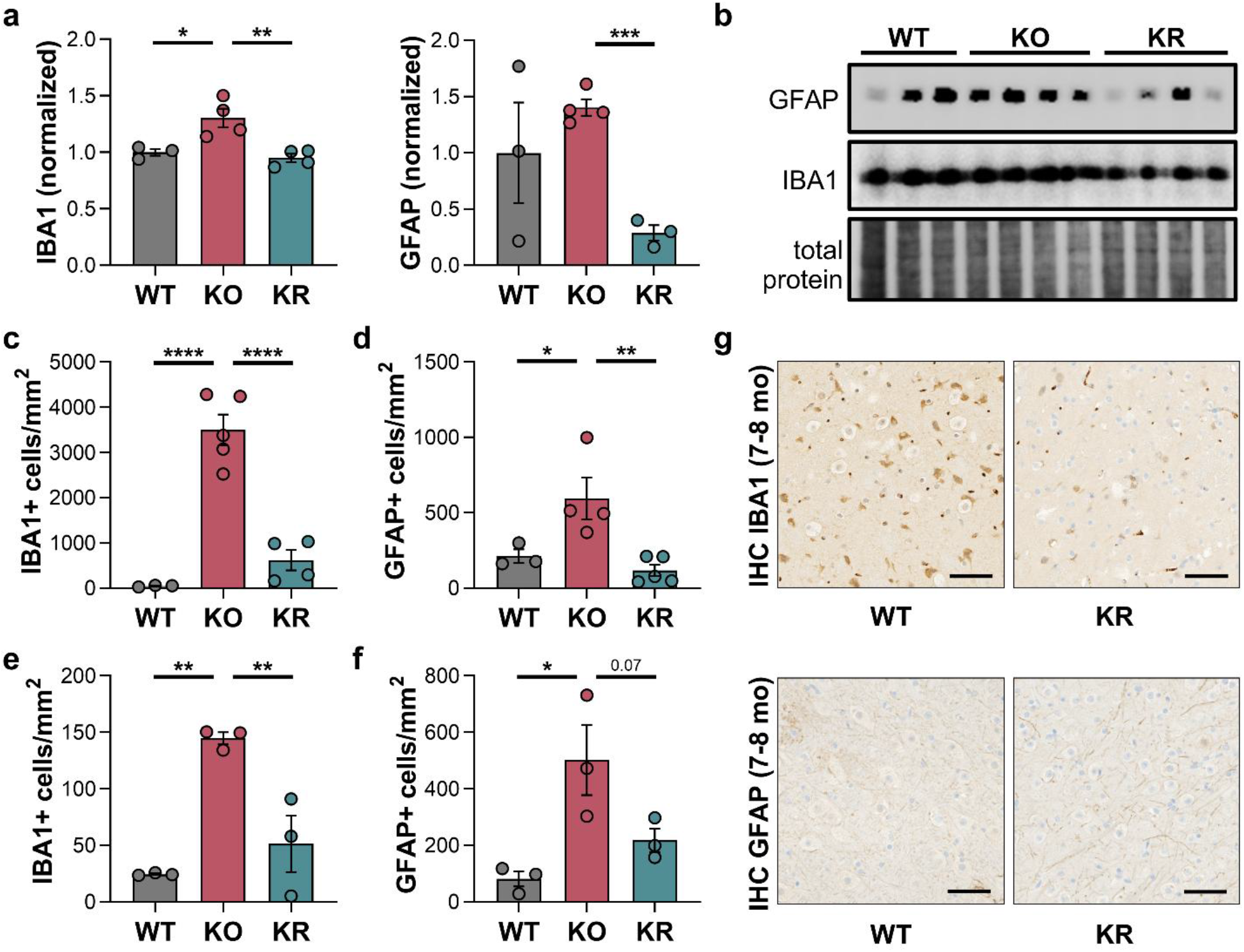
Rapamycin reduces neuroinflammation. **a**, Densitometry of WBs (relative to loading control and normalized to WT levels) of GFAP and IBA1 in whole brain extracts from WT, KO, or KR mice at PND30. **b**, Representative WB images of GFAP and IBA1 in whole brain extracts from WT, KO, or KR mice at PND30. Each lane represents an individual mouse. **c**, Quantification of IBA1+ cells from immunofluorescence staining of thalamus. **d**, Quantification of GFAP+ cells from immunofluorescence staining of thalamus. **e**, Quantification of IBA1+ cells from immunohistochemistry staining of thalamus. **f**, Quantification of GFAP+ cells from immunohistochemistry staining of thalamus. **g**, Representative immunohistochemistry imaging of GFAP and IBA1 in WT and KR mice at 7-8 months. Unless noted otherwise, markers represent individual mice and results in charts are shown as mean ± SEM and statistical analysis was performed using a one-way ANOVA with Šídák multiple comparisons test. *p<0.05, **p<0.01, ***p<0.001, ****p<0.0001, ns=not significant

**Extended Data Figure 5.**
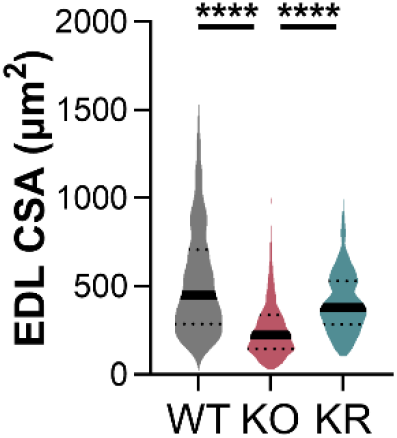
Rapamycin rescues muscle pathology. Analysis of EDL myofiber cross-sectional area in WT, KO, or KR mice at PND20. Statistical analysis of CSA (n=258-1853 fibers counted) was performed using a non-parametric Kruskal-Wallis test with Dunn’s multiple comparisons test.

**Extended Data Figure 6.**
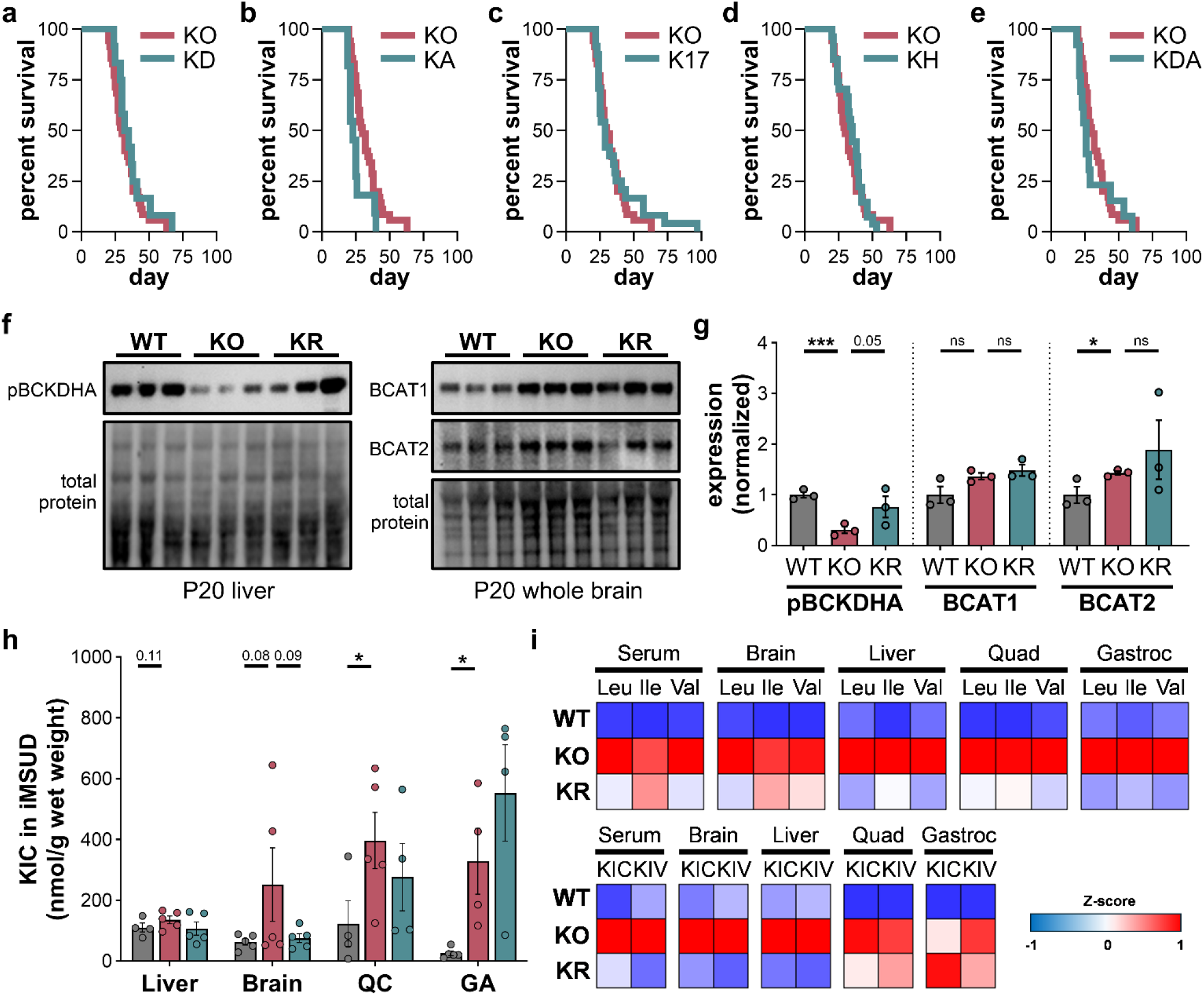
Mechanism of Rescue of Rapamycin in iMSUD. **a**, Survival curves of iMSUD mice treated with vehicle (KO, n=35, red) and DMKG (KD, n=12, teal) starting at PND10 (300 mg/kg/day, i.p. injection). p<0.7889, log-rank with Holm-Šídák correction. **b**, Survival curves of iMSUD mice treated with vehicle (KO, n=35, red) and AMTP (KA, n=11, teal) starting at PND21 (25 mg/kg/day, i.p. injection). p<0.0764, log-rank with Holm-Šídák correction. **c**, Survival curves of iMSUD mice treated with vehicle (KO, n=35, red) and 17-AAG (K17, n=24, teal) starting at PND17 (10 mg/kg/day, i.p. injection). p<0.8449, log-rank with Holm-Šídák correction. **d**, Survival curves of iMSUD mice under normoxia (KO, n=35, red) and hypoxia (KH, n=27, 11% O2, teal) starting at PND25. p<0.8449, log-rank with Holm-Šídák correction. **e**, Survival curves of iMSUD mice treated with vehicle (KO, n=35, red) and dactolisib (KDA, n=13, teal) starting at PND15 (5 mg/kg/day, i.p. injection). p<0.8343, log-rank with Holm-Šídák correction. **f**, Representative western blot (WB) images of phosphorylated BCKDHA in liver extracts, or BCAT1 and BCAT2 in brain extracts from WT, KO, or KR iMSUD mice at PND20. Each individual lane represents the liver or brain from an individual mouse. **g**, Densitometry of phosphorylated BCKDHA, BCAT1, and BCAT2 relative loading control. **h**, Absolute concentrations of the BCKA, KIC, in WT (grey), KO (red) or KR (teal) iMSUD mouse tissues by HPLC after OPD derivatization. **i**, Heat maps of z-scores obtained from the BCAAs leucine, valine, and isoleucine, and BCKAs KIC and KIV in serum, liver, brain, quadriceps, and gastrocnemius in WT, KO, and KR mice at PND20. Unless noted otherwise, markers represent individual mice and results in charts are shown as mean ± SEM and statistical analysis was performed using a one-tailed student’s t-test with Bonferroni correction based on literature precedent for increased BCAAs and BCKAs in MSUD models. *p<0.05, ***p<0.001, ns=not significant

